# Molecular structure of human KATP in complex with ATP and ADP

**DOI:** 10.1101/199034

**Authors:** Kenneth Pak Kin Lee, Jue Chen, Roderick MacKinnon

## Abstract

In many excitable cells KATP channels respond to intracellular adenosine nucleotides: ATP inhibits while ADP activates. We present two structures of the human pancreatic KATP channel, containing the ABC transporter SUR1 and the inward-rectifier K^+^ channel Kir6.2, in the presence of Mg^2+^ and nucleotides. These structures, referred to as quatrefoil and propeller forms, were determined by single-particle cryo-EM at 3.9 Å and 5.6 Å, respectively. In both forms ATP occupies the inhibitory site in Kir6.2. The nucleotide-binding domains of SUR1 are dimerized with Mg^2+^-ATP in the degenerate site and Mg^2+^-ADP in the consensus site. A lasso extension forms an interface between SUR1 and Kir6.2 adjacent to the ATP site in the propeller form, and is disrupted in the quatrefoil form. These structures support the role of SUR1 as an ADP sensor and highlight the lasso extension as a key regulatory element in ADP’s ability to override ATP inhibition.

## Introduction

In 1983, Akinori Noma made the first recordings of ATP-dependent potassium channels in rat cardiac myocytes (Noma, 1983). In membrane patches excised from cardiac myocytes, ATP-dependent K^+^-channels spontaneously open in electrolyte solutions devoid of nucleotides. When exposed to milimolar concentrations of intracellular ATP, these channels become quiescent. Once ATP is removed, the potassium currents re-emerge, thus ATP facilitates channel closure. KATP channels, as they are known, have since been discovered in other cells, including pancreatic beta cells (Cook and Hales, 1984), skeletal (Spruce et al., 1985) and smooth muscle (Standen et al., 1989) fibers and neurons (Karschin et al., 1997; Nelson et al., 2015). Through their selectivity for K^+^ ions, KATP channels return membrane voltage toward the resting (Nernst K^+^) potential (Hibino et al., 2010; Hille, 2001). In addition to ATP, other co-factors also regulate KATP. Mg^2+^-ADP overrides ATP inhibition (Dunne and Petersen, 1986; Kakei et al., 1986; Larsson et al., 1993; Nichols et al., 1996), and phosphatidyl-inositol diphosphate (PIP_2_) is required for activity (Baukrowitz et al., 1998; Hilgemann and Ball, 1996; Shyng and Nichols, 1998). By virtue of their sensitivity to both ATP and ADP, these channels are thought to couple cellular metabolism to membrane excitability.

The physiological importance of KATP channels is underscored by mutations associated with heritable human diseases including congenital hyperinsulinism (Pinney et al., 2008; Saint-Martin et al., 2011; Sharma et al., 2000) and permanent neonatal diabetes mellitus (Aittoniemi et al., 2009; F. M. Ashcroft et al., 2017; Letha et al., 2007). Furthermore, drugs used in the treatment of diabetes mellitis, hypertension and alopecia target KATP channels (Aziz et al., 2014; Feldman, 1985; Rubaiy, 2016; Shorter et al., 2008; Standen et al., 1989; Sturgess et al., 1985).

A broad and rich body of literature based on decades of study describes the biochemical and functional properties of KATP channels (Aguilar-Bryan and Bryan, 1999; Aguilar-Bryan et al., 1998; F. M. Ashcroft et al., 2017; S. J. Ashcroft and F. M. Ashcroft, 1992; Hibino et al., 2010; Nichols, 2016; 2006). These channels were found to be large oligomeric complexes composed of four sulphonylurea receptor (SUR) subunits, which belong to the ABCC family of ABC transporters, and four Kir6 subunits, which are members of the inward rectifier potassium channel family (Clement et al., 1997; Inagaki et al., 1997; Shyng and Nichols, 1997). This peculiar combination of an ABC transporter and an ion channel has been the focus of extensive study and speculation.

Recent cryo-EM structures of a hamster SUR1-mouse Kir6.2 hybrid complex presented the architecture of KATP, revealing a central K^+^ channel surrounded by four SURs (Li et al., 2017; Martin et al., 2017a; 2017b). The SURs extended away from the channel, akin to propellers, and their nucleotide-binding domains (NBDs) were empty and dissociated from each other. In this study we analyze human KATP with adenosine nucleotides bound to SUR1.

## Results and Discussion

### Two structures of human KATP

In humans there are two SUR genes (SUR1 and SUR2) and two Kir6 genes (Kir6.2 and Kir6.1) (Aguilar-Bryan et al., 1995; Babenko et al., 1998; Chutkow et al., 1996; Inagaki et al., 1995a; 1995b; 1995c). The focus of this study is the human KATP channel composed of SUR1 and Kir6.2, which are found in pancreatic beta cells where they play a major role in regulating insulin secretion. To facilitate large-scale expression and purification of the human SUR1-Kir6.2 complex, we fused the C-terminus of SUR1 to the N-terminus of Kir6.2 using a six amino acid linker containing three repeats of the Ser-Ala dyad. This strategy allows the production of human SUR1-human Kir6.2 complex as a tetramer of the fusion construct. This and other peptide linkers ranging from 6 to 14 residues have been used to generate KATP channel fusions that reproduce the electrophysiological and pharmacological characteristics of wildtype octameric KATP channels, as assessed by inhibition from ATP and sulphonylurea as well as activation by ADP and diazoxide (Chan et al., 2008; Clement et al., 1997; Mikhailov et al., 2005; 1998). We also confirmed functional expression of our fusion construct (KATP_em_) in electrophysiology recordings and ATPase measurements (Figure 1-figure supplement 1). For structural studies, KATP_em_ was purified in PMAL-C8 and mixed with Mg^2+^, ATP, vanadate, and C8-PIP_2_ before imaging.

Cryo-EM reconstructions revealed two major conformations (Figure 1A). One is similar to the published structure (“propeller” form) with the exception that the NBDs are closed in our structure (Figure 1C). The other conformation, which we refer to as the quatrefoil form, is dramatically different (Figure 1B). The NBDs are also closed in the quatrefoil form, however, the SUR1 subunits reside in a different location relative to Kir6 (Figures 1B and 1C). Upon further classification and refinement, the propeller form was determined to 5.6 Å resolution and the quatrefoil form was determined to 3.9 Å resolution.

**Figure 1.**
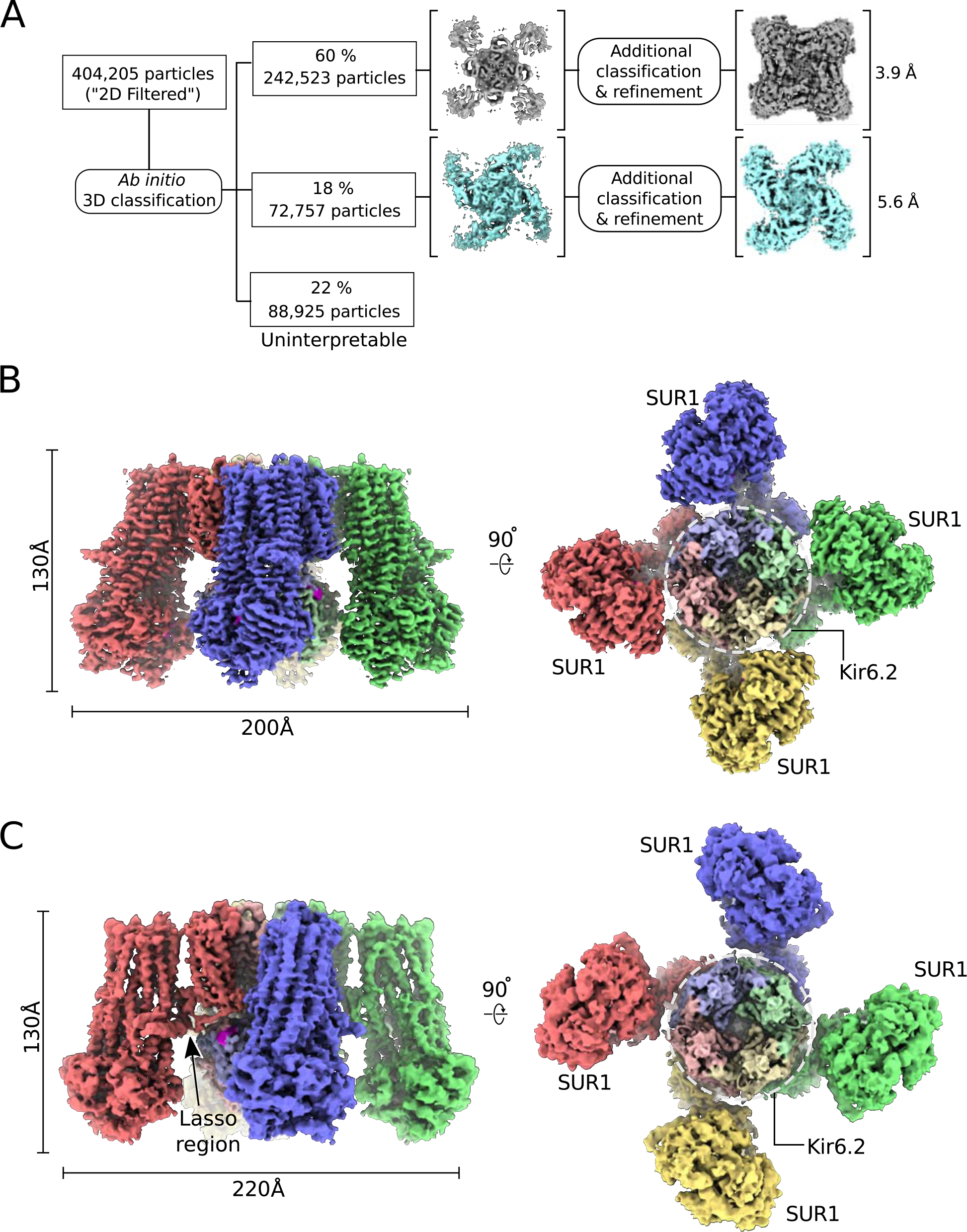
CryoEM reconstruction of the human KATP channel. (**A**) Ab-initio 3D classification separates KATP_em_ particles into two distinct structures - the quatrefoil form (grey) and the propeller form (light blue). (**B**) EM density map of the KATP channel quatrefoil form. Left - Sideview of the EM density map. Right - cytoplasmic view of the EM density map. The CTDs of KIR6.2 is visible in this orientation. SUR1-KIR6.2 fusion protomers are colored red, green, blue or yellow, respectively. The final symmetrized composite map is shown. (**C**) EM density map of the KATP channel propeller form. See also Figure 1-figure supplements 1 to 5.

The EM map of the quatrefoil form was further improved by focused classification and symmetry expansion as implemented in RELION (see methods)(Scheres, 2016). Using this approach, we obtained local maps with substantially enhanced resolution, from which we pieced together a composite 3D image of the KATP_em_ particle (Figure 1B). This enabled building of 1596 out of 1977 residues in the complex and assignment of the amino acid register (Figure 2). In addition, the map was of sufficient quality to identify ligands including Mg^2+^, ATP, and ADP.

**Figure 2.**
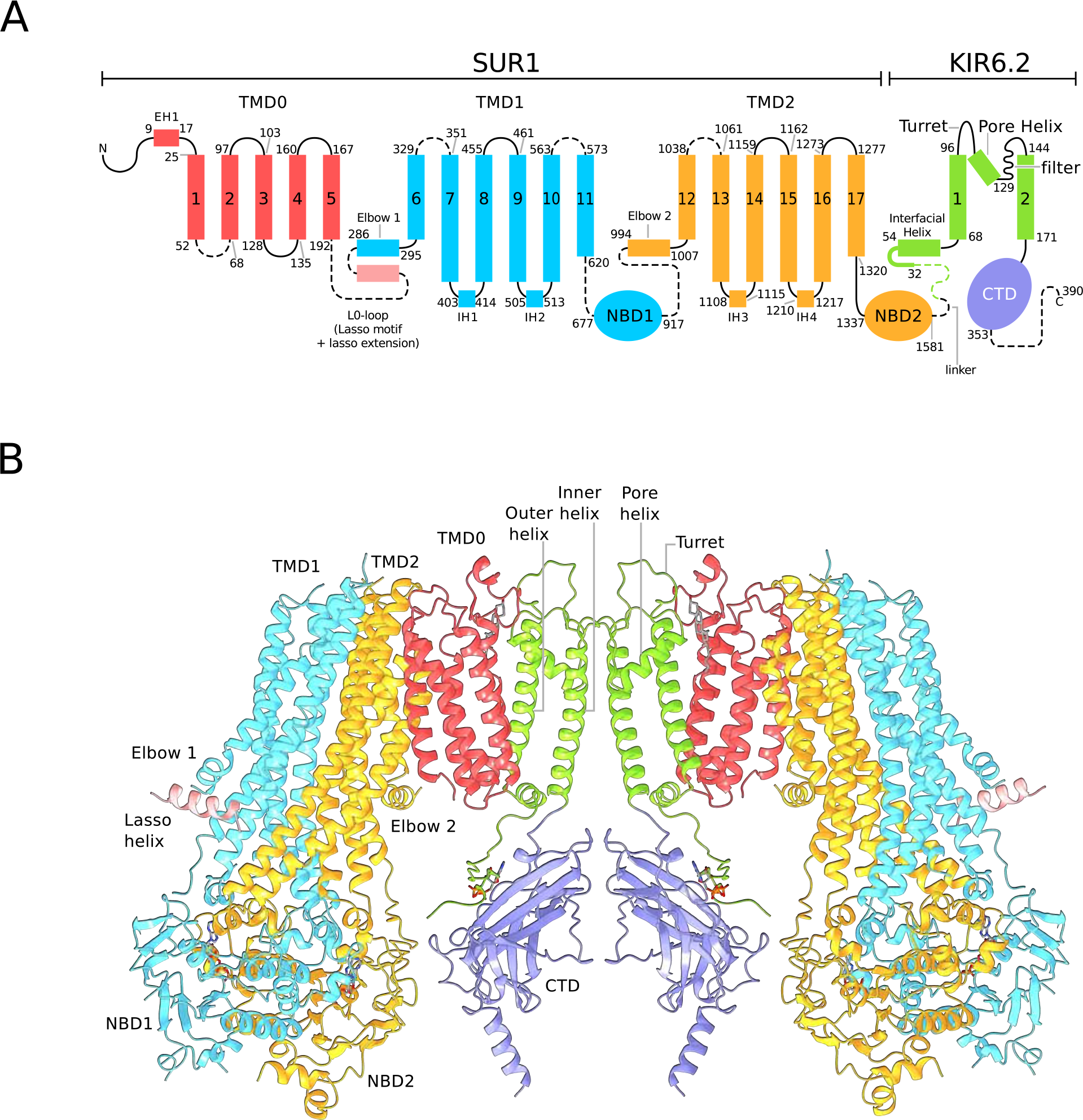
Architecture of the KATP channel. (**A**) Domain architecture of the SUR1-KIR6.2 fusion construct with a flexible six amino-acid linker joins the C-terminal end of SUR1 to the KIR6.2 N-terminus. The numbering scheme conforms to the native human SUR1 and human KIR6.2 sequences. Dashed lines denote regions where clear density was not observed in the quatrefoil form reconstruction. The lasso motif and lasso extension is visible in the propeller form but not the quatrefoil. (**B**) Ribbons representation of two SUR1-KIR6.2 fusion protomers in the quatrefoil form.

In the quatrefoil form KATP is a symmetrical tetramer; each protomer consists of one K^+^ channel subunit and one ABC transporter. There are five domains in the ABC transporter, TMD0, TMD1, NBD1, TMD2 and NBD2 (Figure 2A). The latter four domains form the transporter module. The K^+^ channel consists of a transmembrane domain and a cytoplasmic domain (CTD), which form the ion pathway through the complex (Figure 2B). At the molecular center, the four subunits of Kir6.2 form a canonical inward-rectifier K^+^ channel structure (Figures 1B and 2B). The TMD0s are bound to the channel and hold the four ABC transporters akin to the leaves of a quatrefoil. EM density of the entire complex is very well defined, but residues 193-266, known as the L0-loop, containing a conserved lasso motif (Johnson and J. Chen, 2017; Li et al., 2017; Martin et al., 2017b; Zhang and J. Chen, 2016), is not visible (Figure 2).

A focused 3D classification and refinement strategy coupled with symmetry expansion was also employed to improve the propeller form reconstruction. Whereas the final propeller form EM map is of lower resolution than that of quatrefoil form, secondary structural elements are well defined (Figure 1C). Each domain of the KATP maintains the same structure in both quatrefoil and propeller forms. The major difference between the two structures lies in the different positions of the transporter module relative to the molecular center (Figures 1B and 1C). In addition, the L0-loop is clearly visible in the propeller form albeit at lower resolution (Figure 1C).

We further note that the distance between the C-terminus of SUR1 and the first visible residue of Kir6.2 (Arg32) in the propeller form is 64.2 Å, which is even greater than the corresponding distance in a recent propeller KATP structure without bound Mg^2+^-nucleotides (59.4 Å, PDB ID: 5TWV)(Martin et al., 2017b). Assuming the Cα-Cα distance of adjacent residues in an extended strand spans 3.5 Å (Berg et al., 2002), the maximum distance spanned by the disorder region connecting the C-terminus of SUR1 and Kir6.2 Arg32 (37 a.a. total, including the (SA)_3_ linker) is 118.4 Å, which exceeds the observed distances mentioned above by ~2-fold. Thus, we do not expect the six-amino acid linker used in the KATP_em_ construct to limit the range of motion sampled by SUR1 and Kir6.2 or introduce unnatural distortions to the propeller or quatrefoil forms.

### The nucleotide bound state of SUR1

The transporter module of SUR1 resembles a canonical ABC transporter, TMD1 and TMD2 are domain-swapped, with TM9-10 from TMD1 and TM15-16 from TMD2 reaching across the interface between half-transporters to pack against the neighboring TMD (Figure 3A). The NBDs dimerize in a head-to-tail fashion and nucleotides occupy both ATPase active sites at the dimer interface.

**Figure 3.**
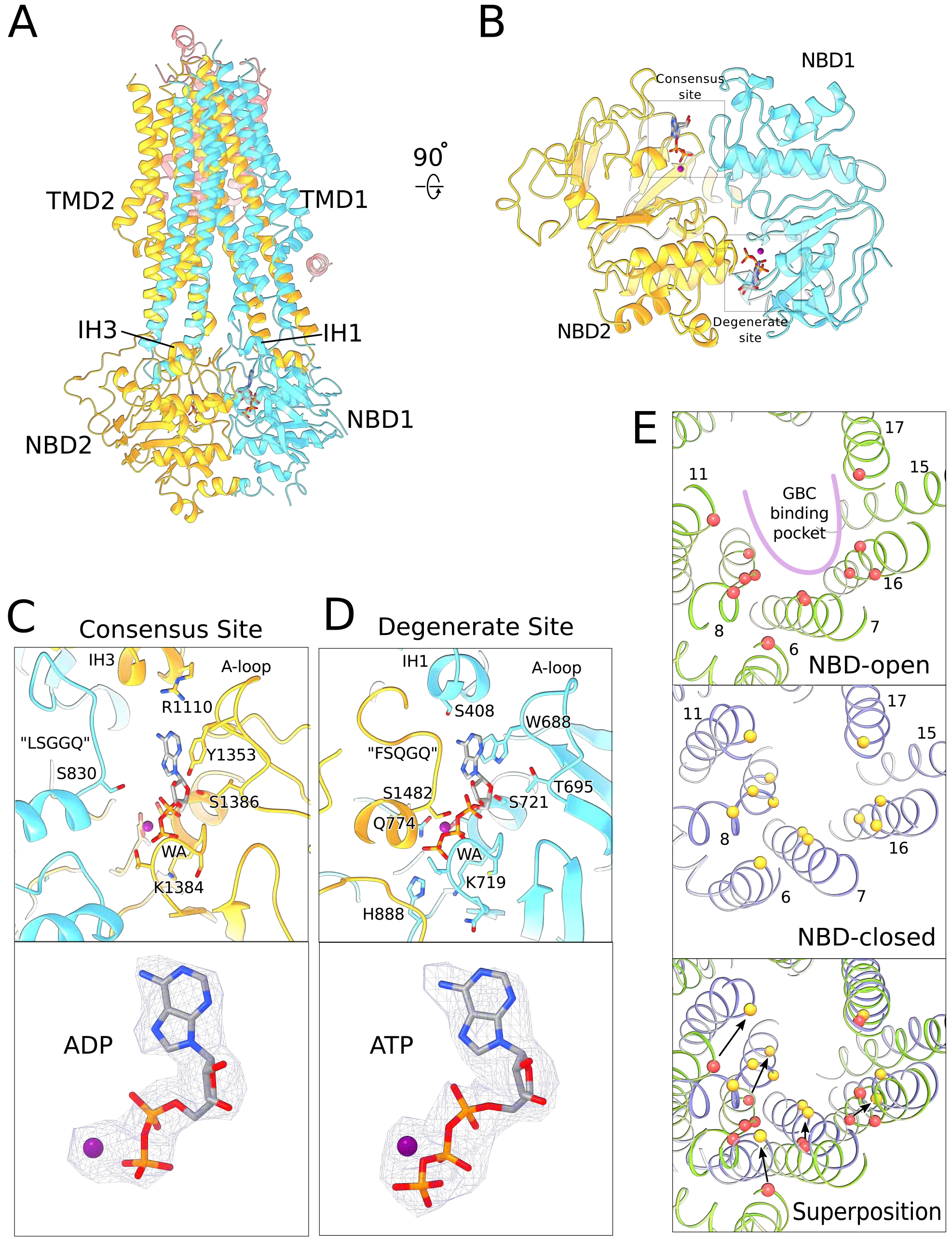
Structure of human SUR1 in complex with Mg^2+^-ATP and Mg^2+^-ADP. (**A**) Ribbons representation of the SUR1 subunit. Color scheme: TMD0, pink; TMD1-NBD1, blue; TMD2-NBD2, yellow. Mg^2+^-ADP and Mg^2+^-ATP are shown in stick model. (**B**) The NBD dimer viewed from the membrane. Mg^2+^-ADP and Mg^2+^-ATP bind to the consensus and degenerate ATPase sites, respectively, to generate an asymmetric NBD dimer. (**C**) Close-up view of the SUR1 consensus ATPase site. Top - Residues in contact with Mg^2+^-ADP are shown. The signature “LSGGQ” motif in NBD1 is disengaged from the bound nucleotide, resulting in a relatively “open” consensus ATPase site. Bottom - shows EM density corresponding to Mg^2+^-ADP as a blue mesh. (**D**) Close-up view of the SUR1 degenerate ATPase site. Top - Residues in contact with Mg^2+^-ATP are shown. The signature sequence in NBD2 is mutated to “FSQGQ” and makes direct contacts with the bound ATP molecule. This results in a “closed” degenerate ATPase site. Bottom - shows EM density corresponding to Mg^2+^-ATP as a blue mesh. (**E**) Structural changes in the glibenclamide (GBC) binding pocket in the NBD-open and NBD-closed states. Top panel - The GBC binding pocket mapped to the NBD-open SUR1 structure (PDB: 5TWV). Residues in contact the GBC are indicated by red spheres (Martin et. al. 2017b). Middle panel - The GBC binding residues (yellow spheres) mapped onto the NBD-closed SUR1 in the quatrefoil form. Bottom panel - Superposition of the NBD-open and NBD-closed states of SUR1 showing the displacement of GBC binding residues upon nucleotide binding. The superposition shown was obtained by alignment of the half-transporter from the NBD-closed and NBD-open SUR1 comprised of: TM9,10,12-14,17 and NBD2. Unless stated otherwise, the quatrefoil form is displayed in all panels.

In an ABC transporter that functions through the principle of alternating access (Jardetzky, 1966), a translocation pathway opens to the extracellular space upon NBD dimerization (Dawson and Locher, 2007). By contrast, in the NBD-dimerized SUR1 the cavity inside the TMDs that would ordinarily form a substrate-binding locus is unusually small. The small cavity is closed off from the extracellular milieu and remains accessible from the cytoplasm (Figure 3A). These features are consistent with SUR1 being a regulator of a K^+^ channel rather than a *bona fide* transporter itself. This notion is further supported by the observation that no substrate has been identified for SUR1 hitherto (Aittoniemi et al., 2009; Schwappach et al., 2000s). Whether the transporter transitions to the outside-open conformation as part of the KATP gating cycle awaits further investigation.

Another unusual feature of SUR1 is the asymmetric configuration of the NBD dimer. Mg^2+^-ATP is bound in the “closed” catalytically inactive degenerate site and Mg^2+^-ADP is bound in the “open” catalytically competent consensus ATPase site (Figures 3B-D). To our knowledge this is the only ABC transporter observed with both Mg^2+^-ADP and Mg^2+^-ATP bound in its NBDs. Since ADP was not supplied to the sample, it either originated from ATP binding followed by hydrolysis or from direct binding of trace ADP present in solution. Given the length of time required to prepare protein samples for cryo-EM analysis it is highly likely that ADP is at equilibrium with its binding site, which would mean the two pathways of occurrence - from ATP hydrolysis within the site versus binding of free ADP - are thermodynamically equivalent. In other words the consensus site in SUR1 binds ADP with higher affinity than ATP and its occupancy by ADP reflects the concentration of free ADP in solution. We will return to the significance of this observation and its consistency with prior studies (Dunne and Petersen, 1986; Dunne et al., 1988; Kakei et al., 1986; Nichols et al., 1996; Tantama et al., 2013) when we consider the mechanism KATP’s ability to sense the metabolic state of a cell.

SUR1 is the target of sulphonylureas that inhibit KATP to promote insulin secretion (Aguilar-Bryan et al., 1995; F. M. Ashcroft et al., 1987; Bryan et al., 2005; Dean and Matthews, 1968; Sturgess et al., 1985). In a recent structure of the hamster-mouse KATP hybrid, the glibenclamide binding site was shown to reside within the TMDs of the inward-facing (i.e. NBD open) transporter module (Martin et al., 2017a) (Figure 3E). In the NBD-dimerized form, residues that comprise the glibenclamide binding pocket move closer towards each other, making the cavity too small to accommodate glibenclamide. This observations is entirely consistent with the hypothesis that glibenclamide stabilizes the inward-facing conformation(Martin et al., 2017a). It functions as a wedge to prevent adenosine nucleotide-mediated closure and thus prevents SUR-mediated regulation of channel activity.

### The inhibitory ATP binding site of human Kir6.2

EM density for the Kir6.2 channel is shown in Figure 4A to illustrate the quality of the map in this region of the transporter-channel complex. Excellent side chain density is observed in both the pore and CTD, which have allowed us to build most of the channel except for disordered flexible regions in the N-and C-termini of Kir6.2 (Figure 4B).

**Figure 4.**
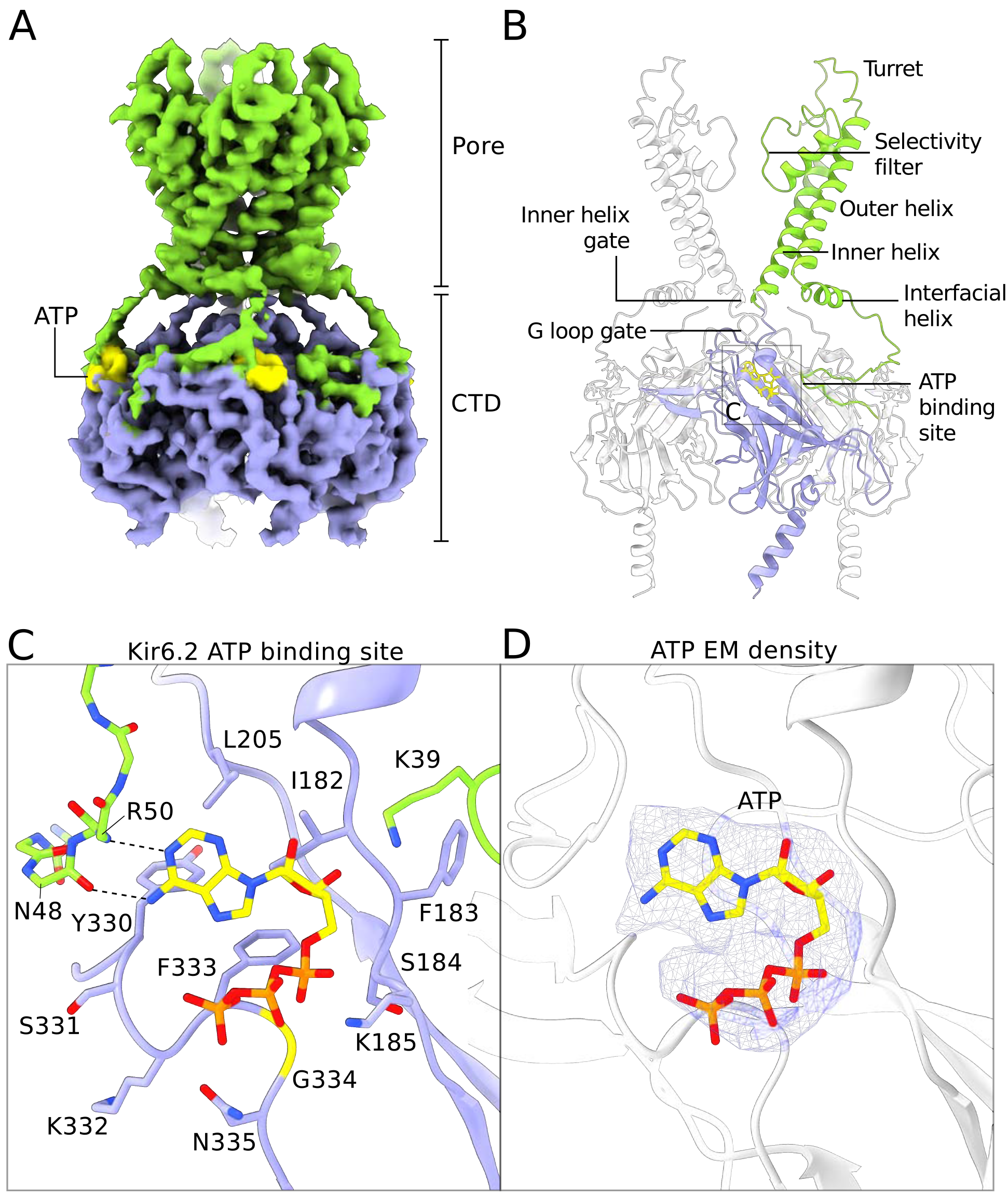
Structure of human Kir6.2 bound to ATP. (**A**) EM density of the Kir6.2 tetramer illustrating the quality of the reconstruction in this region. Sidechain density is clearly observed and β-strands are well resolved. EM densities corresponding to bound ATP molecules are colored yellow. (**B**) Ribbons representation of the Kir6.2 atomic model. For clarity only two pore domains are shown. The CTD domain in the back is also omitted. One Kir6.2 subunit is colored according to the coloring scheme using in Figure 2A. Important structural elements are indicated. The ATP molecule bound to the colored Kir6.2 subunit is shown as a yellow ball-and-stick model. (**C**) Close-up view of the Kir6.2 ATP binding site. Residues in contact with the bound ATP molecule are shown. The N-terminal extension of the interfacial helix from a neighboring Kir6.2 subunit contacts the purine base of ATP via mainchain interactions and is shown in ball-and-stick representation. Hydrogen-bonding interactions between mainchain atoms in the Kir6.2 N-terminus and the adenine base are shown as dashed lines. ATP is shown as a ball-and-stick model, with carbon atoms colored yellow. (**D**) EM density corresponding to the bound ATP molecule, shown as a blue mesh. ATP is displayed in the same fashion as in (**C**). The unusual horseshoe-shaped conformation of ATP is evident. The quatrefoil form is displayed in all panels.

The overall structure of human Kir6.2 is that of an inward-rectifier K^+^-channel with a large cytoplasmic domain. In this structure both the inner helix gate and the G-loop gates are closed (Figure 4B). No PIP_2_ density was observed in the putative PIP_2_ binding sites even though C8-PIP_2_ was present in the sample. By contrast PIP_2_ was present in Kir2 and GIRK inward rectifiers at similar concentrations (Hansen et al., 2011; Whorton and MacKinnon, 2011). Its absence in the KATP structure suggests that ATP, which is known to negatively regulate PIP_2_-mediated activation, may have prevented PIP_2_ binding (Baukrowitz et al., 1998; Shyng and Nichols, 1998).

Four ATP molecules are associated with the Kir6.2 channel (one ATP per subunit); each binds in a shallow pocket on the surface of the CTD (Figure 4). Density for ATP is unambiguous (Figure 4D). Residues N48 and R50 from a neighboring subunit make two hydrogen bonds with the Watson-Crick edge of the adenine base. These interactions likely account for the specific recognition of ATP versus GTP (Schwanstecher et al., 1994; Tucker et al., 1998)(Figure 4C). This same principle of nucleotide selectivity is observed in protein kinases in which main chain atoms of an interdomain linker decode the Watson-Crick edge of the incoming ATP co-substrate (Knighton et al., 1991). Out of the 14 residues that constitute the ATP binding site, mutation of seven are associated with diabetes mellitus, underscoring the physiological importance of ATP regulation (Edghill et al., 2010; Gloyn et al., 2004; Light, 2010; Proks et al., 2004). Notably, G334 and N335 are uniquely found in the Kir6 members of the inward rectifier family. In GIRK the residue analogous to G334 is replaced by histidine, which likely prevents ATP binding by steric hindrance (Masia et al., 2007). These features help to explain why among inward rectifiers Kir6 is uniquely sensitive to ATP.

### ATP binding site is a nexus for communication between regulatory ligands

Three naturally-occurring ligands control the activity of KATP channels: PIP_2_ is required for activity (Hilgemann and Ball, 1996), ATP inhibits activity (Inagaki et al., 1995a; Noma, 1983; Rorsman and Trube, 1985), and ADP potentiates activity (Dunne and Petersen, 1986). All three of these ligands act in a dependent fashion. ATP in functional experiments appears to negatively regulate the action of PIP_2_ (Baukrowitz et al., 1998; Shyng and Nichols, 1998) and ADP appears to override ATP inhibition (Dunne and Petersen, 1986; Kakei et al., 1986; Larsson et al., 1993; Nichols et al., 1996). We observe a structural interconnectedness of the ATP binding site, PIP_2_ binding site and SUR - which is the seat of ADP binding - that seems relevant to their functional dependence.

One side of the inhibitory ATP binding site on Kir6.2 is formed by the N-terminal polypeptide segment that leads to the interfacial “slide” helix (Figures 4C and 5A). A comparison of Kir6.2 and GIRK (Whorton and MacKinnon, 2011) shows that these structural elements are shifted in Kir6.2, presumably to accommodate binding of ATP as residues N48 and R50 reside in this region (Figures 4C and 5A). The shift compresses the PIP_2_ binding site in Kir6.2 compared to GIRK. This compression, if it occurs dynamically when ATP binds, offers a plausible explanation for ATP inhibition through competition with the essential ligand PIP_2_.

**Figure 5.**
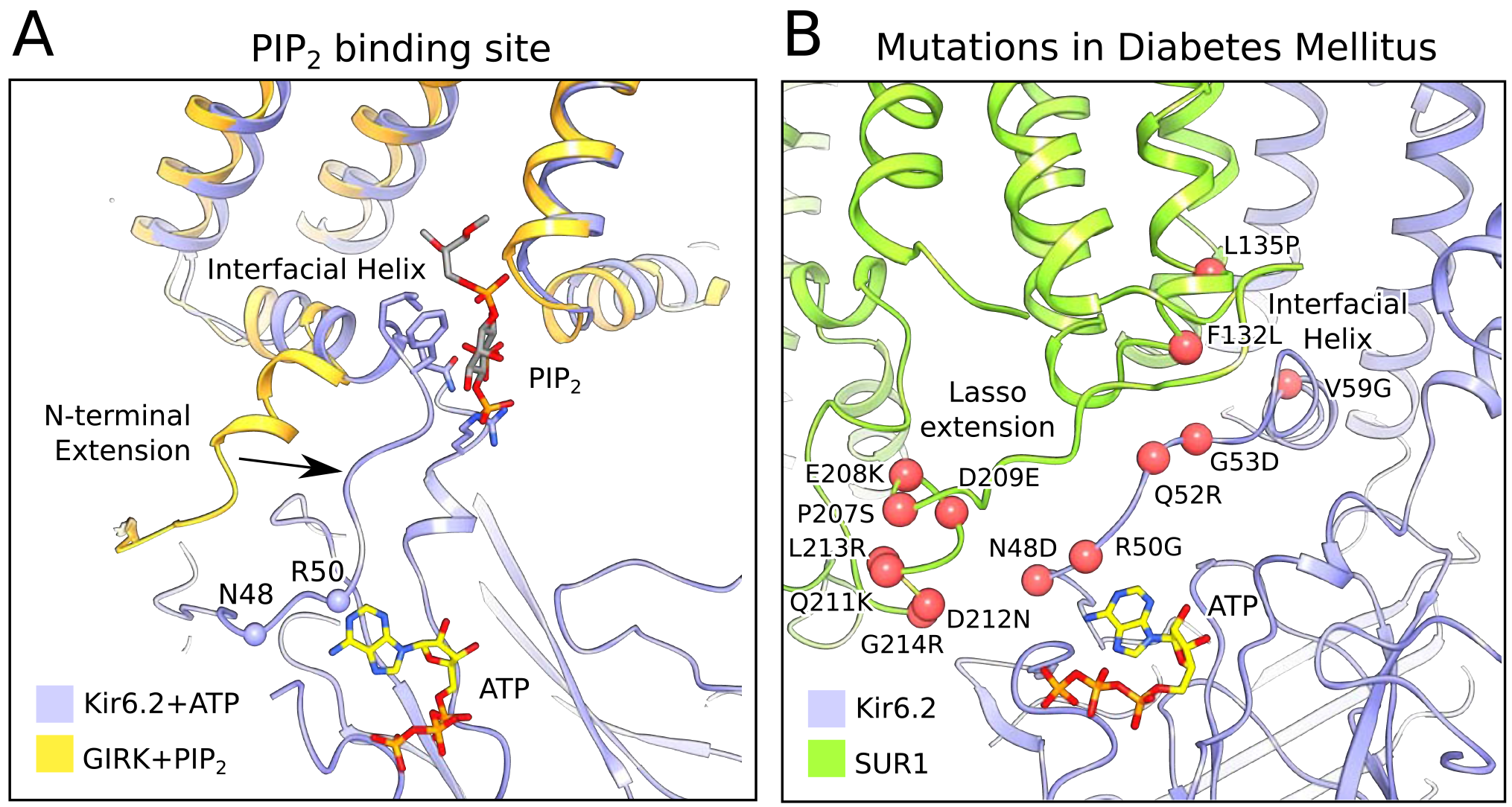
The PIP_2_ binding site and mutations associated with neonatal diabetes. (**A**) Superposition of the ATP-bound Kir6.2 with PIP_2_-bound GIRK (PDB: 3SYA), showing the linkage between the two ligand binding sites. The arrow indicates the movement of the N-terminal extension from the interfacial helix, as compared to GIRK, that accompanies ATP binding. Structural overlay was obtained by alignment of the selectivity-filter sequences of GIRK and Kir6.2. The quatrefoil form of KATP_em_ is shown. (**B**) A close-up view of the Lasso extension-Kir6.2 interface. SUR1 and Kir6.2 in the propeller form are shown in ribbons representation and are colored in green and blue, respectively. Mutations associated with diabetes are indicated by red spheres. The model shown corresponds to the propeller form of KATP_em_.

While ATP inhibition is intrinsic to Kir6.2 (Tucker et al., 1997), SUR1 exerts two opposing influences on ATP inhibition. First, the presence of SUR1 (compared to Kir6.2 expressed in its absence) enhances the potency of ATP inhibition (Tucker et al., 1997). Second, SUR1 permits Mg^2+^-ADP to override ATP inhibition (Nichols et al., 1996; Tucker et al., 1997). The structures of the quatrefoil and propeller forms provide clues as to how SUR1 could accomplish these tasks.

In the propeller form the ATP binding site on Kir6.2 is buttressed by the lasso extension, which attaches SUR1 to the CTD. By virtue of the lasso extension’s close proximity to the ATP binding site it might influence ATP binding. This could account for the fact that SUR1 coexpression potentiates ATP inhibition of KATP by a factor more than 10-fold (Tucker et al., 1997). It could also provide the structural pathway through which ADP binding to SUR overcomes ATP inhibition to activate KATP. Functional data support the likelihood of this possibility. The lasso extension and adjacent amino acids within the ATP binding site (i.e. both sides of this interface between Kir6.2 and SUR1) are hotspots for inherited gain-of-function mutations, which cause diabetes owing to over-activity of KATP channels (Edghill et al., 2010; Gloyn et al., 2004; Light, 2010; Proks et al., 2004). Furthermore, when this interface is locked together by a disulfide cross-link, KATP is permanently inhibited (Pratt et al., 2012). While a mechanistic description of how SUR regulates Kir6 is still unknown, the structure of the lasso extension/ATP-binding site interface and its correlation to disease-causing mutations underscores its potential importance.

### Implications for channel regulation

Formation of the quatrefoil form is associated with disruption of the lasso extension/ATP-binding site interface. SUR1 remains attached to Kir6.2 through its TMD0-channel interface, but rotation of the transport module around the central axis of TMD0 (to convert from propeller to quatrefoil forms) can only occur if the lasso extension releases from its contact adjacent to the ATP site on the CTD. We do not know whether disruption of this interface normally occurs during channel gating or whether it is a energetically unfavorable consequence of strain that is mediated through the interface. Our rationale for even considering the latter possibility is based on a difference between the propeller and quatrefoil structures depicted in Figure 6. In the propeller form, in which the lasso extension/Kir6 interface is intact, the SUR1 transporter modules are held with an offset with respect to the hydrophobic membrane plane, as defined by the position of the Kir6.2 channel and TMD0 domains (Figure 6-figure supplement 2). This offset should produce a hydrophobic mismatch. In the quatrefoil form the SUR1 transport modules reside coplanar, that is, in the same hydrophobic plane as defined by the Kir6 and TMD0 elements (Figure 6B). The consequence of holding the transport modules in an energetically unfavorable position (i.e. with a mismatch) in the propeller must be strain exerted on lasso extension/Kir6 interface.

**Figure 6.**
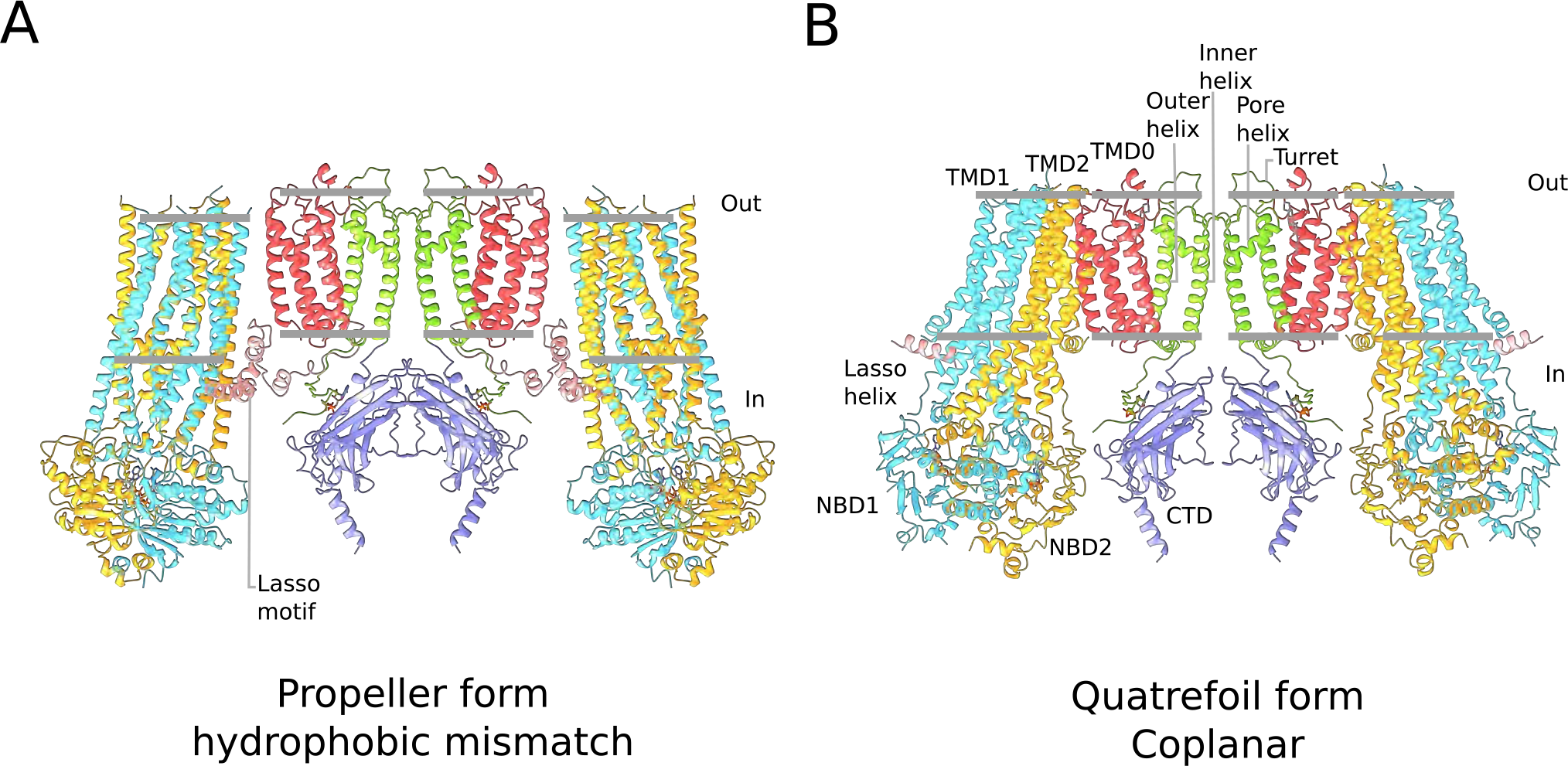
Membrane strain experienced by the quatrefoil and propeller forms of the KATP channel. (**A**) Ribbons representation of the propeller form. Color coding is the same as in Figure 2B. The grey bars indicate the inferred boundaries of the membrane bilayer. The structure of the propeller form predicts a curved membrane surface, which supports that idea that of a strained and thus, higher energy state. (**B**) Ribbons representation of the quatrefoil form, which predicts a flat membrane surface.

When we compare the NBD-dimerized propeller form in this study with the published NBD-open propeller form (Li et al., 2017; Martin et al., 2017b) we observe that dimerization causes a slight tilting of NBD1 away from the center of the complex (Figure 6-figure supplement 3). Thus, dimerization appears to exert force on the lasso extension away from the channel. In this manner nucleotide binding and dimerization of the NBDs could potentially interfere with ATP inhibition. Whether the lasso extension actually dissociates is unknown. The gain-of-function mutations on the lasso extension that should tend to disrupt the interface, and the cross-link that should stabilize it, seem to support the idea that the interface is dynamic (Edghill et al., 2010; Light, 2010; Pratt et al., 2012). Whether or not it is dynamic, we think it likely that dimerization of the NBDs acts through the lasso extension to activate the channel.

The observation of ADP at the SUR1 consensus site is significant. This finding, together with the prior demonstration that channel gating is not coupled to ATP hydrolysis (i.e. gating never appears to violate microscopic reversibility), supports the idea that SUR1 is an ADP sensor (Aittoniemi et al., 2009; Choi et al., 2008; Dunne et al., 1988; Dunne and Petersen, 1986; Gribble et al., 1997; Kakei et al., 1986; Larsson et al., 1993; Proks et al., 2010; Tantama et al., 2013). Because the affinity of the consensus site for ADP appears to be so much higher than for ATP, the channel can potentially sense small changes in ADP levels even in the setting of physiological ATP concentrations. In the quatrefoil form we observe a new interface formed between SUR1 and the Kir6.2 CTD near the ADP binding site (Figure 6-figure supplement 1). Perhaps relevant to this observation, mutation of a single glycine residue to arginine at this interface, found in patients with Congenital Hyperinsulinism completely destroys the stimulatory effect of MgADP in the presence of ATP(de Wet et al., 2012; Stanley et al., 2004). Further study is required to determine whether this interaction is significant to channel gating.

## Methods

### Cell lines

Human embryonic kidney (HEK) cells were cultured in DMEM (GIBCO) medium supplemented with 10% fetal bovine serum at (FBS) 37°C. Sf9 cells were cultured in Sf-900 II SFM medium (GIBCO) at 28°C. HEK293S GnTI‐ cells cultured in Freestyle 293 medium (GIBCO) supplemented with 2% FBS at 37°C.

### Construct design

Synthetic cDNAs encoding human SUR1 and human Kir6.2 were initially cloned into pEG BacMam vectors containing a C-terminal GFP (Goehring et al., 2014). Next we generated a fusion construct in which the C-terminus of SUR1 was linked to the N-terminus of Kir6.2 through a linker consisting 3x (Ser-Ala). To facilitate protein detection and purification, a GFP tag was appended to the C-terminus of KATP_em_ with an intervening PreScission protease cleavage site and cloned into a pEG BacMam vector.

### Electrophysiological recording

Human embryonic kidney (HEK) cells were cultured on coverslips placed in 6-well plates. The cells in each well were transfected with synthetic cDNAs encoding human SUR1, human Kir6.2, or human KATP_em_ cloned into pEG BacMam vectors containing a C-terminal GFP tag using Lipofectamine 3000 (Invitrogen) according to the manufacturer’s instructions. After 36-48hrs, the coverslips were transferred to a recording chamber for patch clamp experiments. The bath solution in the recording chamber contains 5 mM Hepes, pH 7.4, 100 mM NaCl, 2.6 mM CaCl_2_ 1.2 mM MgCl_2_, and 40 mM KCl. Boroscilicate micropipettes were pulled and fire polished. The recording micropipettes were filled with internal solution containing 5 mM Hepes, pH 7.2, 107 mM KCl, 10 mM EGTA, 1.2 mM MgCl_2_, and 1.0 mM CaCl_2_. Pipettes had a resistance of 2-4 MΩ. Whole cell recordings were performed at room temperature using an Axopatch 200B amplifier, a Digidata 1550 digitizer, and pCLAMP software (Molecular Devices). The recordings were low-pass filtered at 1 kHz and sampled at 20 kHz. Cells were kept at a holding potential of −60 mV. Voltage clamp recordings were obtained by consecutively stepping the voltage from −120 mV to +110 mV in 10 mV increments. Cells were returned to the holding potential of −20 mV for 500 ms in between test pulses before the next voltage step.

### Protein expression and purification

Human KATP_em_ was expressed in HEK293S GnTI‐ cells using the BacMam method. In brief, bacmids encoding the Human KATP_em_-GFP fusion were generated using DH10Bac cells according to the manufacturer’s instructions. Bacmam baculoviruses were produced using *Spodoptera frugiperda* Sf9 cells cultured in SF900II SFM medium. For protein expression, suspension cultures of HEK293S GnTI‐ cells cultured in Freestyle 293 medium were infected with bacmam baculovirus at a density of ~3x10^6^ cells/ml. After 24 hrs at 37°C, the infected cultures were supplemented with 10 mM sodium butyrate and grown for a further 48 hrs at 30°C before harvesting. All subsequent manipulations were performed at 4°C.

For large-scale purification, cell pellets from 4 liters of culture were pooled and resuspended in lysis buffer containing 50mM Tris-Cl, pH 8.5, 500mM KCl, 1mM EDTA, 2mM DTT and supplemented with a mixture of protease inhibitors (Tao et al., 2009). The cell suspension was subject to gentle mechanical disruption in a Dounce homogenizer and the resulting lysate was clarified at 39800 x g for 30 min. The crude membrane pellet obtained was resuspended once again in lysis buffer using a few strokes in the Dounce homogenizer and the membrane suspension was stirred for 2 hrs in the presence of 1.5% (w/v) lauryl maltose neopentyl glycol (LMNG) and 0.3% (w/v) cholesteryl hemisuccinate (CHS). The solubilized membranes were clarified by centrifugation at 39800 x g for 45 min and the resulting supernatant was mixed with GFP nanobody-coupled sepharose resin (prepared in-house) by rotation. After 60 min, the resin was collected and washed with 20 column volumes of wash buffer containing 20 mM Tris-Cl, pH 8.5, 300 mM KCl, 1 mM DTT, 1 mM MgCl_2_, 0.5 mM CaCl_2_, 0.1% (w/v) digitonin and 0.01% (w/v) phospholipids (POPC:POPE:POPG =3:1:1; CEG311). Elution of Human KATP_em_ from the GFP nanobody resin was performed by overnight incubation with PreScission protease. The eluted protein was concentrated and then exchanged into amphipols by mixing with PMAL-C8 at amphipol:protein=10:1 (w/w) overnight. The protein:detergent:amphipol mixture was then diluted ten-fold with detergent-free (DF) buffer (20 mM Tris-Cl, pH 8.5, 300 mM KCl) and concentrated once again to a volume of ~500 uL prior to fractionation on a Superose6 column equilibrated with DF buffer (20 mM Tris-Cl, pH 8.5, 300 mM KCl,). Peak fractions were collected and diluted to their final target concentrations before imaging by cryo-EM.

### ATPase activity assay

An NADH-coupled fluorimetric assay was used to measure ATPase activity (Scharschmidt et al., 1979). Mg^2+^-ATP was added to a mixture containing 3.4 µM KATP_em_, 50 mM Tris-Cl, pH 8.0, 300 mM KCl, 30 mM MgCl_2_, 0.5 mM CaCl_2_, 0.1% (w/v) digitonin and 0.01% (w/v) phospholipids (POPC:POPE:POPG =3:1:1; CEG311), 60 µg/mL pyruvate kinase, 32 µg/mL lactate dehydrogenase, 4 mM phosphoenolpyruvate, and 300 µM NADH. Consumption of NADH was measured by monitoring the fluorescence at λ_ex_ = 340 nm and λ_em_ = 445 nm using an Infinite M1000 microplate reader (Tecan). Rates of ATP hydrolysis were calculated by subtracting the rate of fluorescence loss in the absence of KATP_em_ and converting fluorescence loss to nmol NADH per minute using known standards of NADH. Data were then fit by nonlinear regression to the Michaelis-Menten equation to calculate K_M_ and V_max_ values using GraphPad Prism.

### EM data acquisition

Purified human KATP_em_ in DF buffer was diluted to ~0.45 mg/mL using DF buffer and supplemented with 9 mM MgCl_2_, 8mM ATP, 0.5 mM Na_3_VO_4_ and 150 µM C8-PIP_2_. The supplemented protein sample was incubated at room temperature for ~3 hrs and passed through a 0.45 µm filter to remove debris. To prepare cryo-EM grids, 3.5 uL drops of supplemented and filtered human KATP_em_ were applied to glow-discharged Quantifoil R1.2/1.3 400 mesh Au grids. The grids were blotted for 4 s following an incubation period of 15 s at 4°C and 100% humidity before being plunge-frozen in liquid ethane using a Vitrobot Mark IV (FEI). The grids were imaged using a Titan Krios transmission electron microscope (FEI) operated at 300 keV. Automated data collection was performed using SerialEM (Mastronarde, 2003). A K2 Summit direct electron detector (Gatan) was used to record micrographs in super-resolution counting mode with a superresolution pixel size of 0.65 Å. Images were recorded for 10 s over 40 frames using a dose-rate of 8 electrons per pixel per second with a defocus range of 0.9 to 2.5 µm. The total cumulative dose was ~47 electrons per Å^2^ (~1.18 electrons per Å^2^ per frame).

### Image processing and map calculation

Dose-fractionated super-resolution movies were 2x2 down-sampled by Fourier cropping to a final pixel size of 1.3 Å. The down-sampled movie frames were used for grid-based motion correction and dose-filtering with MotionCor2 (Zheng et al., 2017). CTF parameters were estimated from the corrected movie frames using CTFFIND4.1 (Rohou and Grigorieff, 2015). Motion-corrected dose-filtered movie sums were used for interactive and automated particle picking in EMAN2 (Tang et al., 2007). The entire dataset was also manually inspected to eliminate micrographs exhibiting imaging defects including excessive drift, cracked ice or defocus values exceeding the specified range. An initial set of 675,202 particles was obtained from 3,179 images. Particle images were extracted from the motion-corrected dose-filtered images as 300x300 pixel boxes in RELION (Kimanius et al., 2017; Scheres, 2012).

2D classification in RELION was performed to remove spurious images of ice, carbon support and other debris, reducing the particle count to 404,205. We refer to this reduced set of particle images as the “2D filtered” particle set. *Ab initio* 3D classification was performed using stochastic gradient descent and branch-and-bound algorithms implemented in cryoSPARC (Punjani et al., 2017) with C4 symmetry imposed (Figure 1-figure supplement 2). Of the three classes obtained, two classes, containing 60% (Class A, quatrefoil form; 169,766 particles) and 18% (Class B, propeller form; 72,757) of the input particles displayed clear protein features. The remaining class was uninterpretable. The quatrefoil class was refined in cryoSPARC with a reported gold-standard FSC resolution of 3.6 Å but the quality of the density for the SUR1 transporter was rather poor. Refinement of the propeller class in cryoSPARC produced a map with reported resolution of 4.0 Å but it too was characterized by weak density in the SUR1 transporter.

We then tested whether using a different image processing strategy could produce a better result. Using the “2D filtered” particle set as the initial input and the *ab initio* quatrefoil map from cryoSPARC (low-pass filtered to 60 Å) as the starting reference model, we performed iterative cycles of 3D refinement and 3D classification without alignment in RELION. A final round of local refinement was performed in RELION with a soft mask to exclude the amphipol belt surrounding the transmembrane portion of the particle. Over-fitting and over-estimation of resolution introduced by masking was estimated by the high-resolution noise substitution procedure implemented in RELION and the resulting corrected unbiased gold-standard FSC resolution estimates are reported. This procedure resulted in a quatrefoil form reconstruction at FSC=0.143 resolution of 3.9 Å (Class 1; 47282 particles) after masking (Figure 1-figure supplements 2 and 3A).

We observed improved helical density in the transporter module of SUR1 in Class 1 compared to the quatrefoil class from cryoSPARC (Class A). However, the density in the SUR1 NBDs remained weak and poorly resolved, which indicated structural heterogeneity and breakdown of symmetry in that region. Recent advances in statistical image processing methods allow the classification of structural heterogeneity and the recovery of improved local 3D information from inhomogeneous samples exhibiting pseudo-symmetry (Scheres, 2016; Zhou et al., 2015). To improve the Class 1 reconstruction, we divided the cryo-EM density map into eight overlapping sectors for focused 3D classification and focused 3D refinement in RELION (Figure 1-figure supplement 4). The sectors chosen are follows: (1) Kir6.2 CTD tetramer; (2) Kir6.2 channel tetramer; (3) Kir6.2 channel tetramer and four TMD0; (4) Kir6.2 channel tetramer, four TMD0 and one transporter module (TMD1, TMD2, NBD1 and NBD2); (5) two TMD0 and one transporter module; (6) one transporter module; (7) one CTD tetramer and one NBD dimer (NBD1+NBD2); (8) one NBD dimer. Soft masks corresponding to these eight sectors were created using `relion_mask_create` from the RELION package. For sectors 1 to 3, C4 symmetry was enforced during focused 3D classification (without alignment) and focused 3D refinement in RELION. For sectors 4 to 8, the Class 1 particle stack were first artificially expanded 4-fold according to C4 circular symmetry using the RELION command `relion_particle_symmetry_expand` (Scheres, 2016). The symmetry expanded particle stack was then used as input for masked 3D classification and masked 3D refinement in RELION with the focus masks corresponding to sectors 4 to 8. Masked 3D classification was performed without alignment and masked local refinement was performed to optimize alignment parameters. The masked 3D classification and refinement steps were iterated until no further improvement to the reconstructions was observed by manual inspection. For all of the focus maps obtained by this approach, we found significant enhancement to observable features and reported resolution compared to the initial map (Figure 1-figure supplement 3). The quality of the focus maps allowed the identification and modeling bound ligands including ATP, Mg^2+^-ATP and Mg^2+^-ADP as described in the main text. To facilitate map interpretation, we merged focus maps corresponding to all eight sectors into a single composite map using the compositing algorithm implemented in REFMAC5 (Murshudov, 2016). The resultant composite map encompasses the Kir6.2 channel tetramer, four TMD0 and one transporter module, which is equivalent to the volume enclosed in sector 4. The final composite map was sharpened by scaling to the synthetic map calculated from the refined atomic model using `diffmap.exe` (http://grigoriefflab.janelia.org/diffmap). The half-map equivalent of the composite map was prepared according to the procedure described above except half-maps from focused 3D refinements were used in lieu of the full-maps. To generate the symmetrized composite map shown in Figure 1B and Figure 1-figure supplement 3B, the region corresponding to the human KATP_em_ construct (one SUR1 subunit followed by one Kir6.2 subunit) was isolated from the sharpened composite map using UCSF chimera (Pettersen et al., 2004) and symmetrized using `e2proc3d.py` (Tang et al., 2007). Local resolution was estimated using `blocres` with a box size of 18 (Heymann and Belnap, 2007). Representative sections of the cryo-EM density in the quatrefoil form reconstruction are shown in Figure 1-figure supplement 5. Finally, we compared the symmetrized composite map with the map prior to masked 3D refinement and symmetry expansion (Figure 1-figure supplements 3A and 3B). The FSC between the two maps shows a correlation at FSC=0.5 to 6.3 Å indicating that the symmetry expansion, focused refinement and map compositing steps did not introduced global distortions (Figure 1-figure supplement 3C).

We applied a similar focused 3D classification and refinement with symmetry expansion to improve the initial propeller form reconstruction from cryoSPARC (Class B, see Figure 1-figure supplement 2). Briefly, particles corresponding to the propeller form Class B were subjected to additional rounds of *ab initio* 3D classification. A single class corresponding to 16,070 of the starting 72,757 particles displayed improved protein features in the SUR1 portion of the particle. This reconstruction was low-pass filtered to 60 Å and was used as the initial model to refine the stack of 16,070 particles in RELION, which resulted in a reconstruction at FSC=0.143 resolution of 5.6 Å after masking (Figure 1-figure supplement 2). To further improve the propeller reconstruction, iterative focused 3D classification and refinement was performed in RELION using focus masks covering two overlapping sectors: (1) one SUR1 subunit (2) Kir6.2 channel tetramer and four TMD0 domains. Symmetry expansion was performed before focused classification and refinement in Sector 1. Composite maps (full maps and half maps) of the propeller form were prepared according to the procedure described above for the quatrefoil form reconstruction.

### Model building and coordinate refinement

Model building was initially performed in the focus maps corresponding to the quatrefoil form because of their higher quality. The cryo-EM structure of bovine MRP1 (PDB: 5UJA) was used as a reference structure to generate the starting model for building the SUR1 transporter module. Briefly, the MRP1 model (without TMD0) was first mutated to match the SUR1 sequence using CHAINSAW (Stein, 2008) while keeping only side chains of conserved residues. The model was then divided into two pieces, each corresponding to one half transporter (one TMD plus one NBD). The SUR1 half transporters were then docked into EM density separately by rigid-body fitting using the `fitmap` function in UCSF Chimera (Pettersen et al., 2004) followed by manual rebuilding in Coot (Emsley et al., 2010). To build the SUR1 TMD0 domain, the EM density in the focus maps was of sufficient quality to allow *de novo* manual building in Coot. The crystal structure of GIRK (PDB: 3SYA) was used a reference structure to generate the starting model for building the Kir6.2 tetramer. CHAINSAW was again used to mutate the GIRK model to match the Kir6.2 sequence and eliminating non-conserved side chains. The Kir6.2 pore module and CTD were separated and converted to tetramers by applying the appropriate symmetry operations. The pore and CTD tetramer models were then rigid-body fitted independently into EM density in UCSF Chimera to allow for rotation of the CTD relative to the pore. The fitted Kir6.2 models were then used as starting points for manual rebuilding in Coot.

The SUR1 transporter, the TMD0 domain, and the Kir6.2 tetramer channel were first built as independent initial models by consulting all available focus maps. B-factor sharpening was performed locally in Coot “on-the-fly” to optimize observable map features for model building. Side chains were not modeled for residues with poor density. The initial models were then merged into a single consensus model containing the Kir6.2 channel tetramer, four TMD0 domains and one transporter module. Composite maps of the quatrefoil (one full map and two half maps) were then prepared as described above. Prior to automatic real space refinement, one additional round of manual rebuilding in Coot was then performed using the consensus model and the composite full map.

Automatic real space refinement of the consensus model was performed against one of the composite half-maps using `phenix.real_space_refine` (Adams et al., 2010) with 4-fold symmetry imposed for the Kir6.2 channel tetramer and the four TMD0 with the application of NCS constraints. Tight secondary structure and geometric restraints were used to minimize overfitting. Manual rebuilding in Coot was alternated with automated refinement in `phenix.real_space_refine`.

For cross-validation, FSC curves were calculated between refined models and the composite halfmap used for refinement (FSC_work_) or the composite half-map not used at any point during refinement (FSC_free_) (Figure 1-figure supplement 3D). These curves were inspected after each round of automated real-space refinement in `phenix.real_space_refine` to monitor the effects of overfitting.

Regions that did not allow accurate establishment of amino acid register were modeled as a polyalanines. Regions with weak or no density were not modeled and are indicated by dashed lines in Figure 2A. The quality of the final model was evaluated by MolProbity (V. B.Chen et al., 2012) and EMRinger (Barad et al., 2015) (Table 1).

**Table 1.**

| KATP with Mg<sup>2+</sup>-ATP |  |  |  |
| --- | --- | --- | --- |
| Data acquisition |  |  |  |
|  | Microscope | Titan Krios |  |
|  | Voltage (kV) | 300 |  |
|  | Camera | Gatan K2 Summit |  |
|  | Camera mode | Super-resolution |  |
|  | Defocus range (μm) | 0.9 to 2.5 |  |
|  | Pixel size (Å) | 1.3 (super-resolution=0.65) |  |
|  | Movies | 3,179 |  |
|  | Frames/movie | 40 |  |
|  | Total electron dose (e-/Å²) | 47 |  |
|  | Exposure time (s) | 10 |  |
|  | Dose rate (e-/pixel/s) | 8 |  |
|  |  | Quatrefoil form | Propeller form |
| Reconstruction |  |  |  |
|  | Software | RELION | RELION |
|  | Symmetry | C4 | C4 |
|  | Particle number | 47,282 | 16,070 |
|  | Resolution (masked, Å) | 3.9 | 5.6 |
| Model statistics |  |  |  |
|  | Residues built | 1596/1977 | 1667/1977 |
|  | Map CC (masked) | 0.825 | 0.743 |
|  | Cβ outliers | 0 | 0 |
|  | Molprobity score | 2.18 | 2.33 |
|  | EMRinger score | 1.82 | 0.95 |
| Ramachandran |  |  |  |
|  | Favored (%) | 92.48 | 90.99 |
|  | Allowed (%) | 7.51 | 8.73 |
|  | Outliers (%) | 0.00 | 0.28 |
| RMS deviations |  |  |  |
|  | Bond length | 0.007 | 0.006 |
|  | Bond angles | 1.074 | 1.134 |

Model building into the propeller form reconstruction made use of the atomic model built into the higher quality quatrefoil reconstruction. The Kir6.2 pore, the CTD, the TMD0 and transporter module from the quatrefoil form coordinates were independently docked into the propeller form reconstruction by rigid-body fitting. The agreement between the fitted models and the propeller form EM density was excellent and required little additional adjustment. The L0-loop in SUR1, which contains the lasso helix, the lasso motif and the lasso extension was ordered in the propeller form map, in contrast to the quatrefoil form. Given that the sequence of the lasso-motif in SUR1 and MRP1 is well conserved, we reasoned that the structure of this region in SUR1 should be very similar to that observed in MRP1. To build the lasso-motif in the propeller form density, we extracted the coordinates of the lasso-motif from the bMRP1 model (residues 205 to 248) and mutated it to match the SUR1 sequence using CHAINSAW while keeping sidechains of only conserved residues. This initial model (corresponding to residues 215 to 255 in SUR1) was docked into the lasso motif of the propeller form EM map by rigid-body fitting. The resulting model showed good agreement with the lasso density and required little additional manual adjustment in coot. EM density corresponding to bulky sidechains in this region allowed us to confidently establish the amino-acid register of the lasso-motif. Regions with less defined densities were built as polyalanine models. Automated real-space refinement, manual re-building and cross-validation were performed as described above for the quatrefoil form structure. The quality of the final propeller form model was evaluated by MolProbity and EMRinger (Table 1).

All structure figures were generated using UCSF ChimeraX (Goddard et al., 2017).

## Author contributions

K.L. performed the experiments. K.L, J.C. and R.M. designed the experiments, analyzed the results and prepared the manuscript.

## Acknowledgements

We thank M. Ebraham and J. Sortis at the Evelyn Gruss Lipper Cryo-EM Resource Center at Rockefeller University for assistance in data collection, members of the MacKinnon and Chen laboratories for helpful discussions. K.L. was supported by the Human Frontiers Science Program Fellowship. R.M. and J.C. are investigators in the Howard Hughes Medical Institute.

## Author Information

Cryo-EM density maps of the KATP quatrefoil and propeller forms bound to nucleotides have been deposited in the electron microscopy data bank under accession code EMD-XXXX and XXXX, respectively. Atomic coordinates of the KATP quatrefoil and propeller forms bound to nucleotides have been deposited in the protein data bank under accession code XXXX and XXXX.

---

**Figure 1-figure supplement 1.**
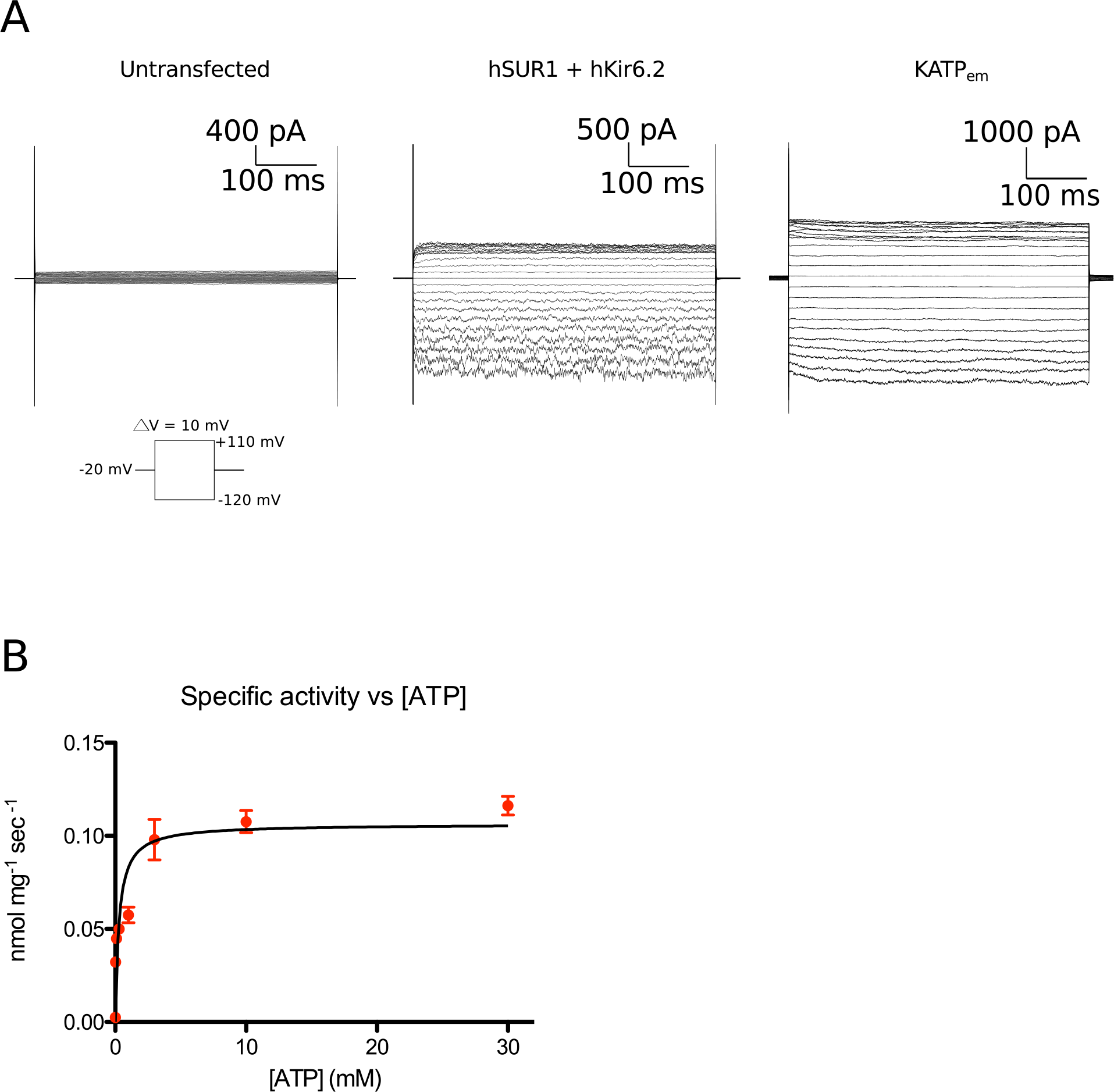
Functional characterization of the human KATP channel. (**A**) Whole cell recordings of human SUR1-KIR6.2 fusion recombinantly expressed in HEK cells. Voltage families stepping between −120mV and +110mV at 10mv intervals were recorded in whole cell mode. Left - untransfected control. Middle - co-transfected human SUR1 and human Kir6.2. Right - transfected with the human SUR1-Kir6.2 fusion used for structure determination. (**B**) ATPase activity of KATP_em_ purified in digitonin. By non-linear regression of the Michaelis-Menten equation, the K_m_ for ATP was determined to be 284 ± 75 µM and the maximal ATPase activity was determined to be 6.4 ± 0.3 nmol/mg/min (assuming the molecular weight of KATP_em_ to be 220 kDa, the specific turnover rate is 1.4 min^−1^.

**Figure 1-figure supplement 2.**
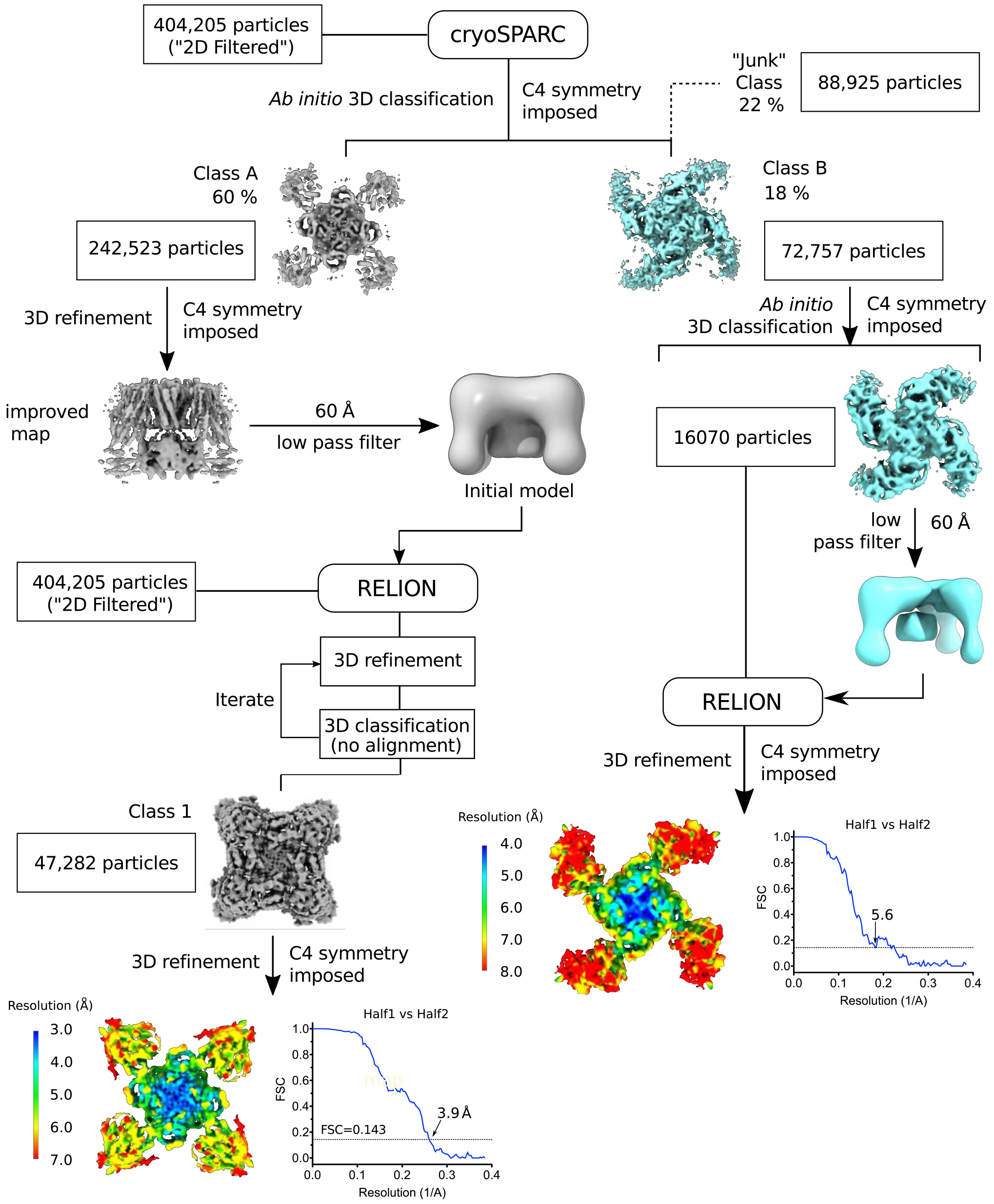
Flow chart of the image processing workflow. The steps leading up to the determination of the quatrefoil and propeller form structures are shown. Symmetry expansion, focused 3D classification, and refinement (see Methods) are omitted for clarity. Local resolution of the quatrefoil and propeller form maps estimated by Blocres are shown at the end of the data processing steps. Gold-standard FSC curves corresponding to the quatrefoil and propeller form are shown next to the mapped local resolution estimates.

**Figure 1-figure supplement 3.**
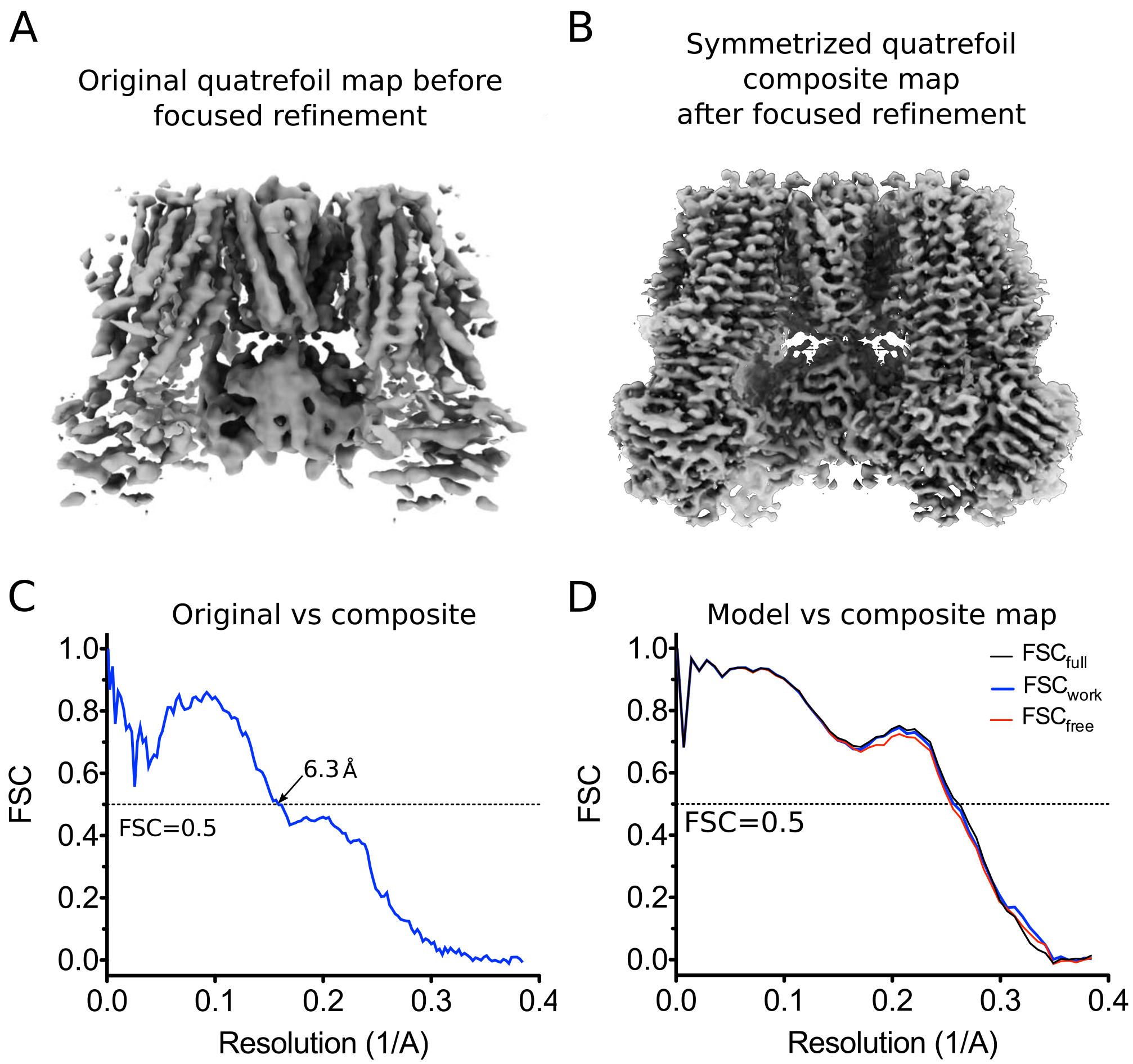
CryoEM reconstruction of the quatrefoil form KATP channel. (**A**) The quatrefoil reconstruction before focused classification and refinement. The map shown is low-pass filtered to 3.9 Å without sharpening. (**B**) The symmetrized quatrefoil form composite map after symmetry expansion, iterative focused 3D classification and refinement. The map shown is low-pass filtered to 3.5 Å and sharpened with a B-factor of −100 Å^2^. (**C**) Fourier shell correlation (FSC) curves between the original quatrefoil map before focused refinement and the symmetrized quatrefoil composite map. (**D**) FSC curves for cross-validation of the quatrefoil form model. The curves show the correlation between a synthetic map calculated from the final model and composite maps of the quatrefoil form reconstruction. Model map vs. Full composite map (FSC_full_, black). Model map vs. half composite map used for structure refinement (FSC_work_, blue). Model map vs. half composite map excluded from structure refinement (FSC_free_, red).

**Figure 1-figure supplement 4.**
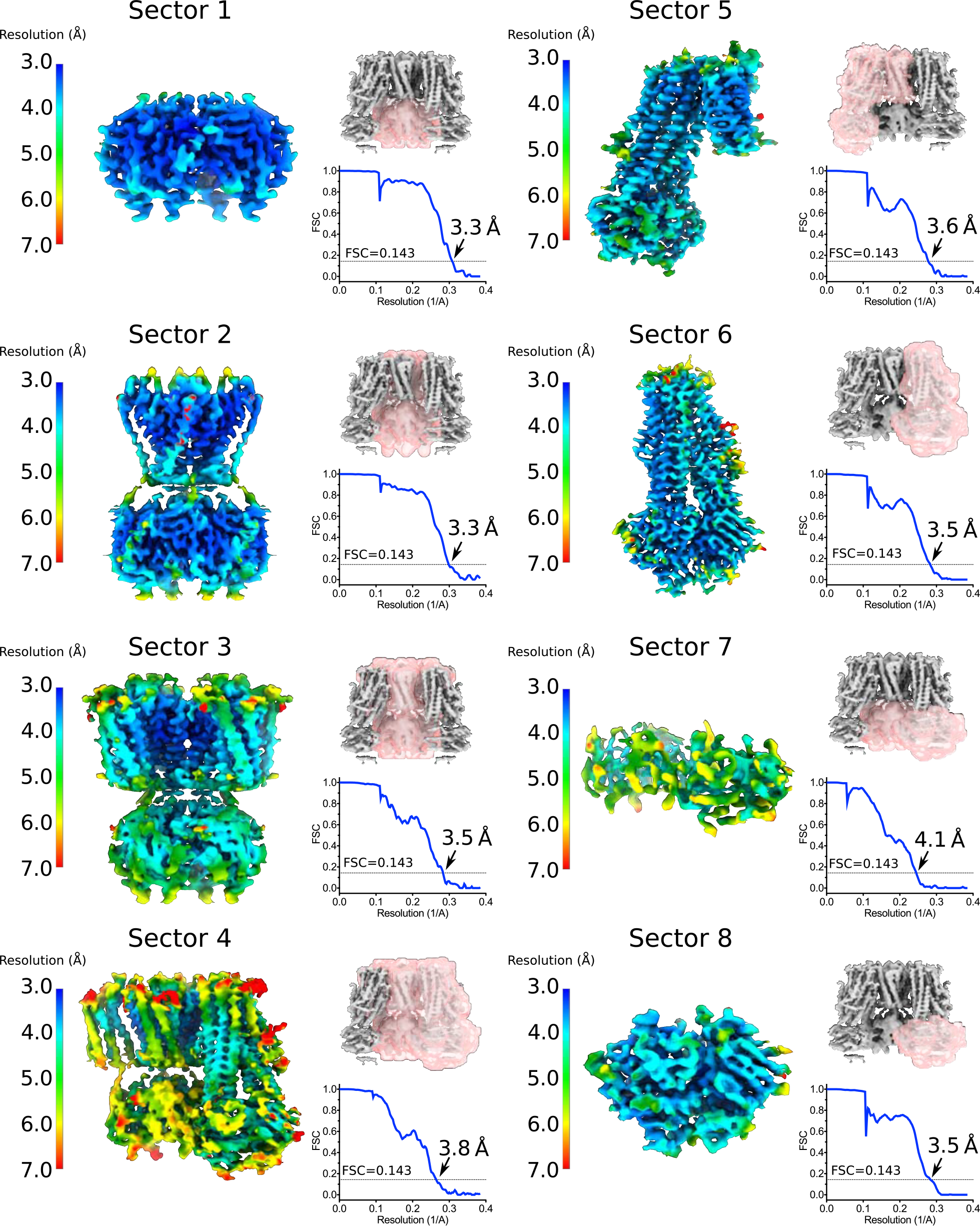
Focus maps of the quatrefoil form KATP channel used for composite map generation. Local reconstructions using various focus masks corresponding to eight overlapping regions (denoted as Sectors) in the quatrefoil form. The maps are the final focus maps obtained after iterative focused 3D classification and refinement, colored according to local resolution estimates from Blocres. Focus masks for each sector are shown as transparent pink surfaces on the initial quatrefoil reconstruction. Also shown are the gold-standard FSC curves for each local reconstruction. The resolution at FSC=0.143 is indicated. Sector 1 - Kir6.2 CTD (tetramer). Sector2 - Kir6.2 channel (tetramer). Sector 3 - Kir6.2 (tetramer) and four TMD0s. Sector 4 - Kir6.2 (tetramer), four TMD0s and one transporter module. Sector 5 - one transporter module and two neighbouring TMD0s. Sector 6 - onne transporter module. Sector 7 - CTD (tetramer) and NBD-dimer from one transporter module. Sector 8 - NBD-dimer.

**Figure 1-figure supplement 5.**
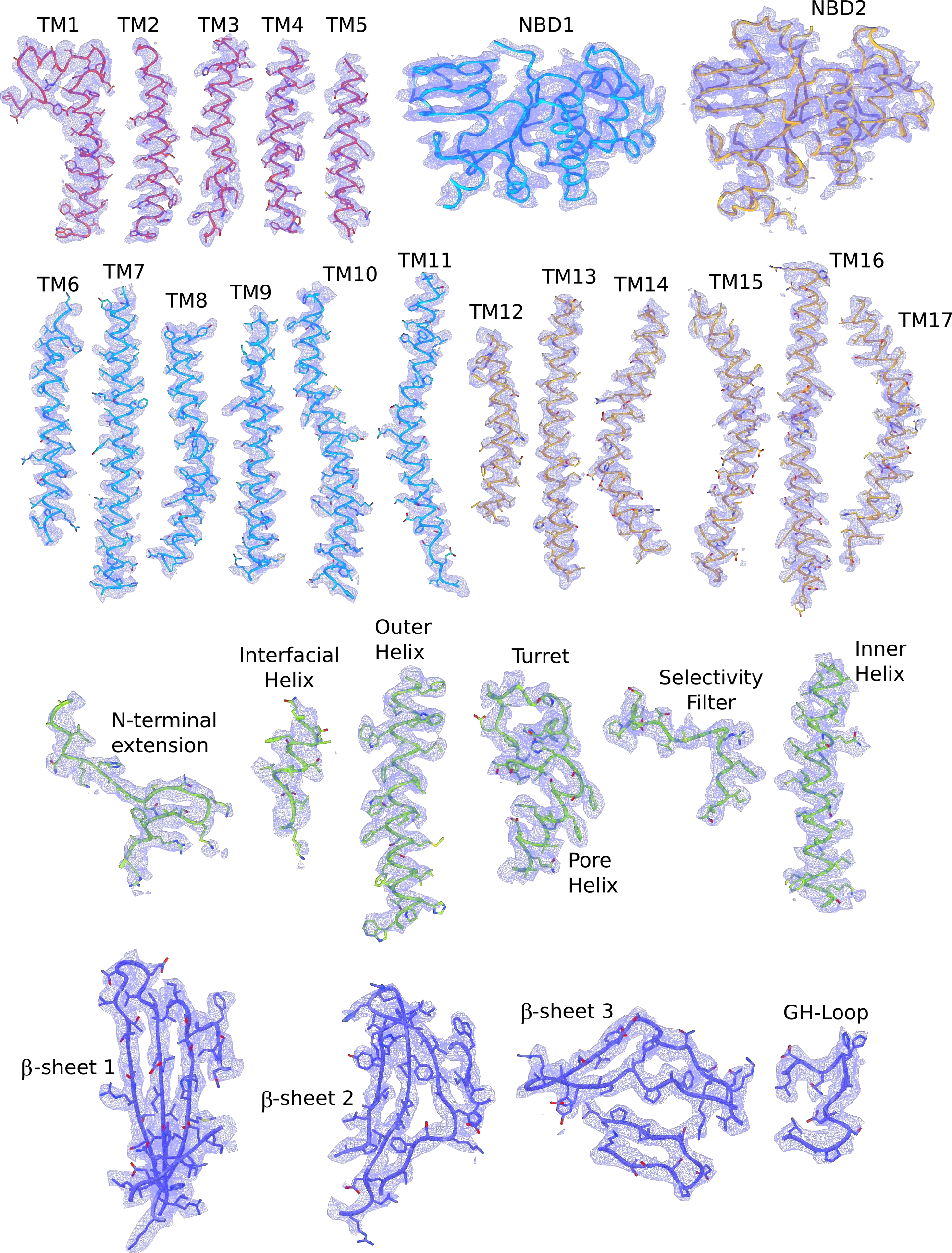
Cryo-EM densities of different regions of the quatrefoil form KATP channel. The atomic model shown are colored according to the color scheme used in Figure 2b (TMD0 - red; TMD 1 and NBD1 - light blue; TMD2 and NBD2 - gold; Kir6.2 pore - green; Kir6.2 CTD - dark blue). Note that the segment corresponding to the Lasso extension and Lasso motif was not built because of poor density in that region. In contrast, these regions are ordered and visible in the propeller form reconstruction correspond to the composite map of the quatrefoil form sharpened with a B-factor of −100 Å^2^.

**Figure 6-figure supplement 1.**
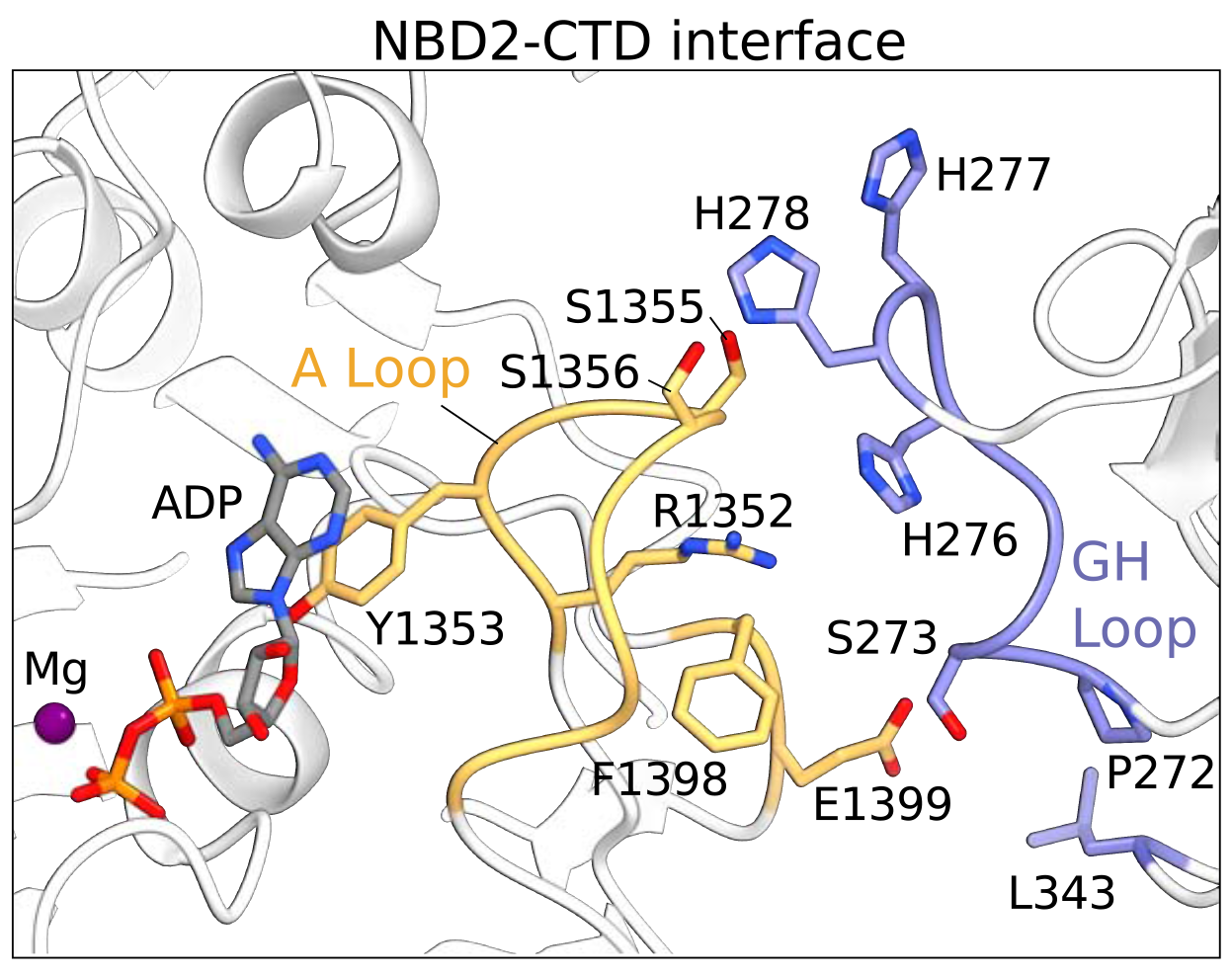
The NBD2-CTD interface. The NBD2 of SUR1 and the CTD of KIR6.2 are shown in ribbons representation. Residues populating the NBD2-CTD interface reside mainly in the A-loop of NBD2 in SUR1 (orange) and GH-loop of the CTD in KIR6.2 (blue). The sidechains of residues contributing to the interface are shown. Mg^2+^-ADP in the consensus ATPase site of SUR1 is immediately adjacent to the NBD2-CTD interface and is shown as a ball-and-stick model.

**Figure 6-figure supplement 2.**
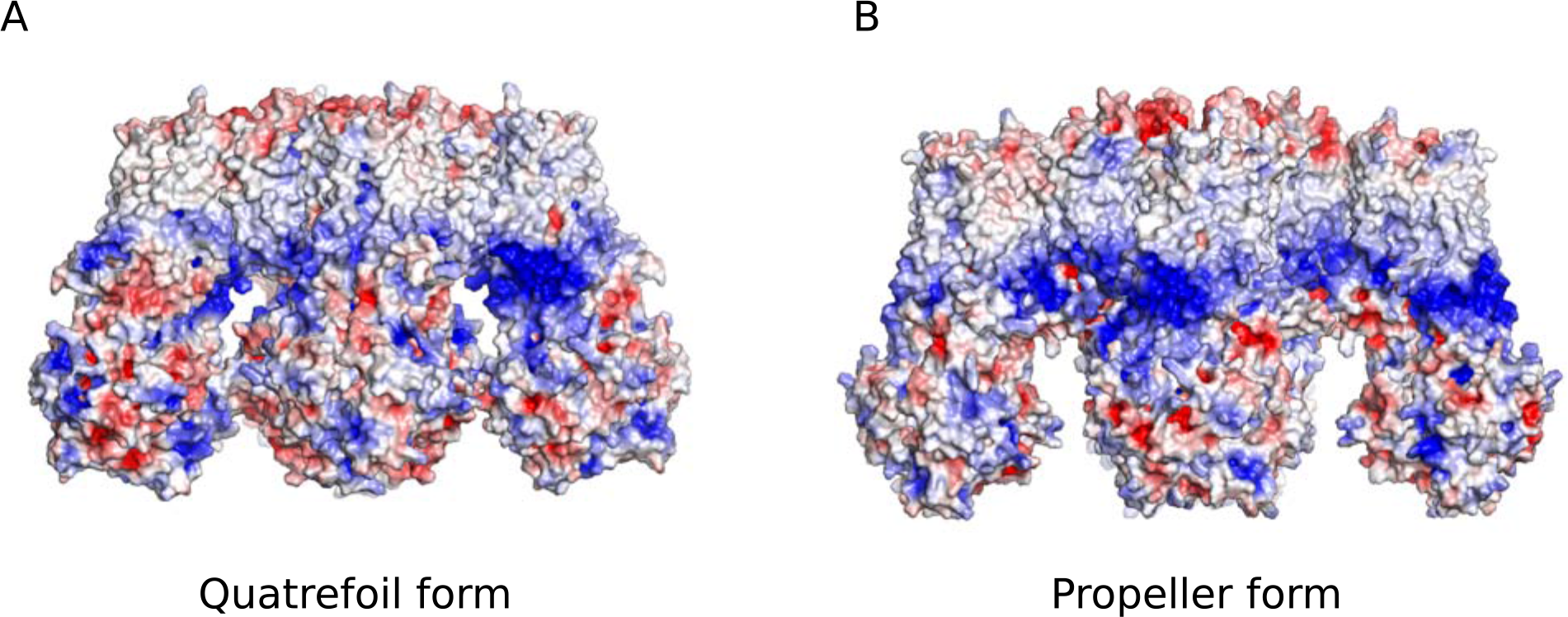
Comparison of the electrostatic potential surfaces of the quatrefoil and propeller forms. Side-views of the electrostatic potential surfaces of both quatrefoil and propeller forms are shown to illustrate the offset in the hydrophobic membrane delimited regions between the two forms. (**A**) Electrostatic surface representation of the quatrefoil form. (**B**) The propeller form in the same representation.

**Figure 6-figure supplement 3.**
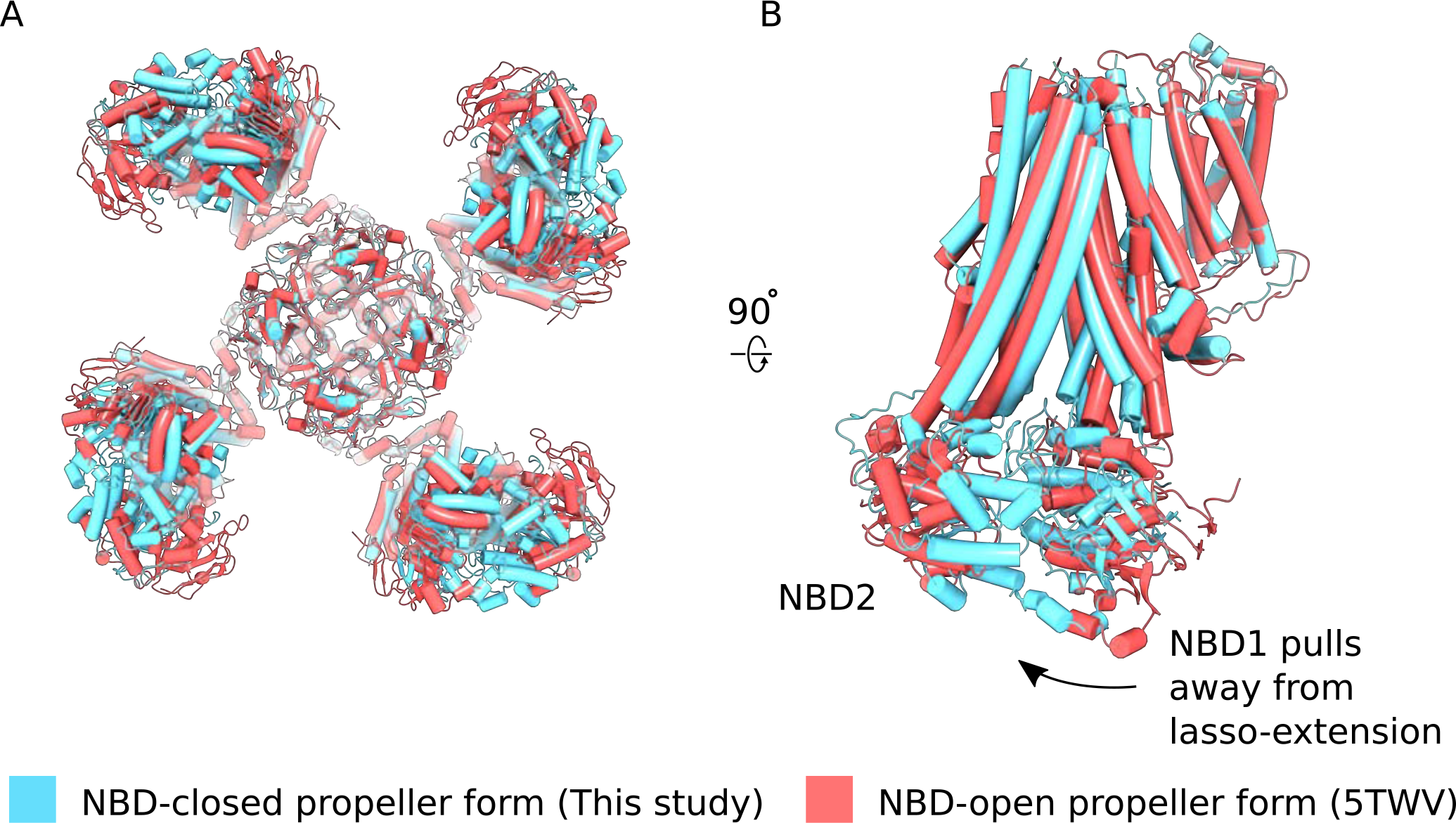
Comparison of NBD-closed and NBD-open propeller forms. (**A**) The propeller forms of KATP bound to Mg^2+^-ATP and Mg^2+^ADP NBD-closed state (blue) or Mg^2+^-nucleotide free NBD-open state (PDB ID: 5TWV) (red) brought into co-incidence by alignment of their selectivity filters. The structure alignment is viewed from the bottom. The two propeller form states are very similar overall but differences in the disposition of NBDs of the two states are evident. (**B**) Side-view of one SUR1 subunit in the NBD-closed and NBD-open states from the alignment shown in (**A**). Nucleotide engagement correlates with tilting of NBD1 away from the lasso-extension. See main text for discussion.

## Reference

Adams, P. D., Afonine, P. V., Bunkóczi, G., Chen, V. B., Davis, I. W., Echols, N., Headd, J. J., Hung, L.-W., Kapral, G. J., Grosse-Kunstleve, R. W., McCoy, A. J., Moriarty, N. W., Oeffner, R., Read, R. J., Richardson, D. C., Richardson, J. S., Terwilliger, T. C., Zwart, P. H., 2010. PHENIX: a comprehensive Python-based system for macromolecular structure solution. Acta Crystallogr. D Biol. Crystallogr. 66: 213–221. doi:10.1107/S0907444909052925

Aguilar-Bryan, L., Bryan, J., 1999. Molecular biology of adenosine triphosphate-sensitive potassium channels. Endocr. Rev. 20: 101–135. doi:10.1210/edrv.20.2.0361

Aguilar-Bryan, L., Clement, J. P., Gonzalez, G., Kunjilwar, K., Babenko, A., Bryan, J., 1998. Toward understanding the assembly and structure of KATP channels. Physiological Reviews 78: 227–245.

Aguilar-Bryan, L., Nichols, C. G., Wechsler, S. W., Clement, J. P., Boyd, A. E., Gonzalez, G., Herrera-Sosa, H., Nguy, K., Bryan, J., Nelson, D. A., 1995. Cloning of the beta cell high-affinity sulfonylurea receptor: a regulator of insulin secretion. Science 268: 423–426.

Aittoniemi, J., Fotinou, C., Craig, T. J., de Wet, H., Proks, P., Ashcroft, F. M., 2009. Review. SUR1: a unique ATP-binding cassette protein that functions as an ion channel regulator. Philos. Trans. R. Soc. Lond B Biol. Sci. 364: 257–267. doi:10.1098/rstb.2008.0142

Ashcroft, F. M., Kakei, M., Kelly, R. P., Sutton, R., 1987. ATP-sensitive K^+^ channels in human isolated pancreatic B-cells. FEBS Letters 215: 9–12.

Ashcroft, F. M., Puljung, M. C., Vedovato, N., 2017. Neonatal Diabetes and the KATP Channel: From Mutation to Therapy. Trends Endocrinol. Metab. 28: 377–387. doi:10.1016/j.tem.2017.02.003

Ashcroft, S. J., Ashcroft, F. M., 1992. The sulfonylurea receptor. Biochim. Biophys. Acta 1175: 45–59.

Aziz, Q., Thomas, A. M., Gomes, J., Ang, R., Sones, W. R., Li, Y., Ng, K.-E., Gee, L., Tinker, A., 2014. The ATP-sensitive potassium channel subunit, Kir6.1, in vascular smooth muscle plays a major role in blood pressure control. Hypertension 64: 523–529. doi:10.1161/HYPERTENSIONAHA.114.03116

Babenko, A. P., Gonzalez, G., Aguilar-Bryan, L., Bryan, J., 1998. Reconstituted human cardiac KATP channels: functional identity with the native channels from the sarcolemma of human ventricular cells. Circ. Res. 83: 1132–1143.

Barad, B. A., Echols, N., Wang, R. Y.-R., Cheng, Y., DiMaio, F., Adams, P. D., Fraser, J. S., 2015. EMRinger: side chain-directed model and map validation for 3D cryo-electron microscopy. Nat. Methods 12: 943–946. doi:10.1038/nmeth.3541

Baukrowitz, T., Schulte, U., Oliver, D., Herlitze, S., Krauter, T., Tucker, S. J., Ruppersberg, J. P., Fakler, B., 1998. PIP_2_ and PIP as determinants for ATP inhibition of KATP channels. Science 282: 1141–1144. doi:10.1126/science.282.5391.1141

Berg, J. M., Tymoczko, J. L., Stryer, L., 2002. Biochemistry, Fifth Edition. W. H. Freeman.

Bryan, J., Crane, A., Vila-Carriles, W. H., Babenko, A. P., Aguilar-Bryan, L., 2005. Insulin secretagogues, sulfonylurea receptors and K(ATP) channels. Curr. Pharm. Des. 11: 2699–2716.

Chan, K. W., Wheeler, A., Csanády, L., 2008. Sulfonylurea receptors type 1 and 2A randomly assemble to form heteromeric KATP channels of mixed subunit composition. J Gen Physiol 131: 43–58. doi:10.1085/jgp.200709894

Chen, V. B., Arendall, W. B. III, Headd, J. J., Keedy, D. A., Immormino, R. M., Kapral, G. J., Murray, L. W., Richardson, J. S., Richardson, D. C., 2012. MolProbity: all-atom structure validation for macromolecular crystallography, in: Crystallography of Biological Macromolecules. International Union of Crystallography, Chester, England, pp. 694–701. doi:10.1107/97809553602060000884

Choi, K.-H., Tantama, M., Licht, S., 2008. Testing for violations of microscopic reversibility in ATP-sensitive potassium channel gating. JPhys Chem B 112: 10314–10321. doi:10.1021/jp712088v

Chutkow, W. A., Simon, M. C., Le Beau, M. M., Burant, C. F., 1996. Cloning, tissue expression, and chromosomal localization of SUR2, the putative drug-binding subunit of cardiac, skeletal muscle, and vascular KATP channels. Diabetes 45: 1439–1445.

Clement, J. P. IV, Kunjilwar, K., Gonzalez, G., Schwanstecher, M., Panten, U., Aguilar-Bryan, L., Bryan, J., 1997. Association and Stoichiometry of KATP Channel Subunits. Neuron 18: 827–838. doi:10.1016/S0896-6273(00)80321-9

Cook, D. L., Hales, N., 1984. Intracellular ATP directly blocks K^+^ channels in pancreatic B-cells. Nature 311: 271–273. doi:10.1038/311271a0

Dawson, R. J. P., Locher, K. P., 2007. Structure of the multidrug ABC transporter Sav1866 from Staphylococcus aureus in complex with AMP-PNP. FEBS Letters 581: 935–938. doi:10.1016/j.febslet.2007.01.073

de Wet, H., Shimomura, K., Aittoniemi, J., Ahmad, N., Lafond, M., Sansom, M. S. P., Ashcroft, F. M., 2012. A universally conserved residue in the SUR1 subunit of the KATP channel is essential for translating nucleotide binding at SUR1 into channel opening. The Journal of Physiology 590: 5025–5036. doi:10.1113/jphysiol.2012.236075

Dean, P. M., Matthews, E. K., 1968. Electrical Activity in Pancreatic Islet Cells. Nature 219: 389–390. doi:10.1038/219389a0

Dunne, M. J., Petersen, O. H., 1986. Intracellular ADP activates K^+^ channels that are inhibited by ATP in an insulin-secreting cell line. FEBS Letters 208: 59–62.

Dunne, M. J., West-Jordan, J. A., Abraham, R. J., Edwards, R. H. T., Petersen, O. H., 1988. The gating of nucleotide-sensitive K^+^ channels in insulin-secreting cells can be modulated by changes in the ratio ATP4-/ADP3-and by nonhydrolyzable derivatives of both ATP and ADP. J. Membrain Biol. 104: 165–177. doi:10.1007/BF01870928

Edghill, E. L., Flanagan, S. E., Ellard, S., 2010. Permanent neonatal diabetes due to activating mutations in ABCC8 and KCNJ11. Rev Endocr Metab Disord 11: 193–198. doi:10.1007/s11154-010-9149-x

Emsley, P., Lohkamp, B., Scott, W. G., Cowtan, K., 2010. Features and development of Coot. Acta Crystallogr. D Biol. Crystallogr. 66: 486–501. doi:10.1107/S0907444910007493

Feldman, J. M., 1985. Glyburide: a second-generation sulfonylurea hypoglycemic agent. History, chemistry, metabolism, pharmacokinetics, clinical use and adverse effects. Pharmacotherapy 5: 43–62.

Gloyn, A. L., Pearson, E. R., Antcliff, J. F., Proks, P., Bruining, G. J., Slingerland, A. S., Howard, N., Srinivasan, S., Silva, J. M.C.L., Molnes, J., Edghill, E. L., Frayling, T. M., Temple, I. K., Mackay, D., Shield, J. P. H., Sumnik, Z., van Rhijn, A., Wales, J. K. H., Clark, P., Gorman, S., Aisenberg, J., Ellard, S., Njølstad, P. R., Ashcroft, F. M., Hattersley, A. T., 2004. Activating mutations in the gene encoding the ATP-sensitive potassium-channel subunit Kir6.2 and permanent neonatal diabetes. N. Engl. J. Med. 350: 1838–1849. doi:10.1056/NEJMoa032922

Goddard, T. D., Huang, C. C., Meng, E. C., Pettersen, E. F., Couch, G. S., Morris, J. H., Ferrin, T. E., 2017. UCSF ChimeraX: Meeting Modern Challenges in Visualization and Analysis. Protein Sci. doi:10.1002/pro.3235

Goehring, A., Lee, C.-H., Wang, K. H., Michel, J. C., Claxton, D. P., Baconguis, I., Althoff, T., Fischer, S., Garcia, K. C., Gouaux, E., 2014. Screening and large-scale expression of membrane proteins in mammalian cells for structural studies. Nat Protoc 9: 2574–2585. doi:10.1038/nprot.2014.173

Gribble, F. M., Tucker, S. J., Ashcroft, F. M., 1997. The essential role of the Walker A motifs of SUR1 in K-ATP channel activation by Mg-ADP and diazoxide. The EMBO Journal 16: 1145–1152. doi:10.1093/emboj/16.6.1145

Hansen, S. B., Tao, X., MacKinnon, R., 2011. Structural basis of PIP_2_ activation of the classical inward rectifier K^+^ channel Kir2.2. Nature 477: 495–498. doi:10.1038/nature10370

Heymann, J. B., Belnap, D. M., 2007. Bsoft: image processing and molecular modeling for electron microscopy. J. Struct. Biol. 157: 3–18. doi:10.1016/j.jsb.2006.06.006

Hibino, H., Inanobe, A., Furutani, K., Murakami, S., Findlay, I., Kurachi, Y., 2010. Inwardly rectifying potassium channels: their structure, function, and physiological roles. Physiological Reviews 90: 291–366. doi:10.1152/physrev.00021.2009

Hilgemann, D. W., Ball, R., 1996. Regulation of cardiac Na+,Ca2+ exchange and KATP potassium channels by PIP_2_. Science 273: 956–959.

Hille, B., 2001. Ion channels of excitable membranes.

Inagaki, N., Gonoi, T., Clement, J. P., Namba, N., Inazawa, J., Gonzalez, G., Aguilar-Bryan, L., Seino, S., Bryan, J., 1995a. Reconstitution of IKATP: an inward rectifier subunit plus the sulfonylurea receptor. Science 270: 1166–1170.

Inagaki, N., Gonoi, T., Seino, S., 1997. Subunit stoichiometry of the pancreatic beta-cell ATP-sensitive K^+^ channel. FEBS Letters 409: 232–236. doi:10.1016/S0014-5793(97)00488-2

Inagaki, N., Inazawa, J., Seino, S., 1995b. cDNA sequence, gene structure, and chromosomal localization of the human ATP-sensitive potassium channel, uKATP-1, gene (KCNJ8). Genomics 30: 102–104. doi:10.1006/geno.1995.0018

Inagaki, N., Tsuura, Y., Namba, N., Masuda, K., Gonoi, T., Horie, M., Seino, Y., Mizuta, M., Seino, S., 1995c. Cloning and functional characterization of a novel ATP-sensitive potassium channel ubiquitously expressed in rat tissues, including pancreatic islets, pituitary, skeletal muscle, and heart. J. Biol. Chem. 270: 5691–5694.

Jardetzky, O., 1966. Simple allosteric model for membrane pumps. Nature 211: 969–970.

Johnson, Z. L., Chen, J., 2017. Structural Basis of Substrate Recognition by the Multidrug Resistance Protein MRP1. Cell 168: 1075–1085.e9. doi:10.1016/j.cell.2017.01.041

Kakei, M., Kelly, R. P., Ashcroft, S. J. H., Ashcroft, F. M., 1986. The ATP-sensitivity of K^+^ channels in rat pancreatic B-cells is modulated by ADP. FEBS Letters 208: 63–66. doi:10.1016/0014-5793(86)81533-2

Karschin, C., Ecke, C., Ashcroft, F. M., Karschin, A., 1997. Overlapping distribution of K(ATP) channel-forming Kir6.2 subunit and the sulfonylurea receptor SUR1 in rodent brain. FEBS Letters 401: 59–64.

Kimanius, D., Forsberg, B., Lindahl, E., 2017. Accelerated Cryo-EM Structure Determination with Parallelisation using GPUs in Relion-2. Biophys. J. 112: 575A–575A.

Knighton, D. R., Zheng, J. H., Eyck, Ten, L. F., Ashford, V. A., Xuong, N. H., Taylor, S. S., Sowadski, J. M., 1991. Crystal structure of the catalytic subunit of cyclic adenosine monophosphate-dependent protein kinase. Science 253: 407–414. doi:10.1126/science.1862342

Larsson, O., Ammälä, C., Bokvist, K., Fredholm, B., Rorsman, P., 1993. Stimulation of the KATP channel by ADP and diazoxide requires nucleotide hydrolysis in mouse pancreatic beta-cells. The Journal of Physiology 463: 349–365. doi:10.1113/jphysiol.1993.sp019598

Letha, S., Mammen, D., Valamparampil, J. J., 2007. Permanent neonatal diabetes due to KCNJ11 gene mutation. The Indian Journal of Pediatrics 74: 947–949. doi:10.1007/s12098-007-0175-y

Li, N., Wu, J.-X., Ding, D., Cheng, J., Gao, N., Chen, L., 2017. Structure of a Pancreatic ATP-Sensitive Potassium Channel. Cell 168: 101–110.e10. doi:10.1016/j.cell.2016.12.028

Light, P., 2010. The molecular mechanisms and pharmacotherapy of ATP-sensitive potassium channel gene mutations underlying neonatal diabetes. PGPM 145–17. doi:10.2147/PGPM.S6969

Martin, G. M., Kandasamy, B., DiMaio, F., Yoshioka, C., Shyng, S.-L., 2017a. Anti-diabetic drug binding site in K ATPchannels revealed by Cryo-EM. bioRxiv 1–33. doi:10.1101/172908

Martin, G. M., Yoshioka, C., Rex, E. A., Fay, J. F., Xie, Q., Whorton, M. R., Chen, J. Z., Shyng, S.-L., 2017b. Cryo-EM structure of the ATP-sensitive potassium channel illuminates mechanisms of assembly and gating. Elife 6: 213. doi:10.7554/eLife.24149

Masia, R., Koster, J. C., Tumini, S., Chiarelli, F., Colombo, C., Nichols, C. G., Barbetti, F., 2007. An ATP-Binding Mutation (G334D) in KCNJ11 Is Associated With a Sulfonylurea-Insensitive Form of Developmental Delay, Epilepsy, and Neonatal Diabetes. Diabetes 56: 328–336. doi:10.2337/db06-1275

Mastronarde, D. N., 2003. SerialEM: A Program for Automated Tilt Series Acquisition on Tecnai Microscopes Using Prediction of Specimen Position. Microscopy and Microanalysis 9: 1182–1183. doi:10.1017/S1431927603445911

Mikhailov, M. V., Campbell, J. D., de Wet, H., Shimomura, K., Zadek, B., Collins, R. F., Sansom, M. S. P., Ford, R. C., Ashcroft, F. M., 2005. 3-D structural and functional characterization of the purified KATP channel complex Kir6.2-SUR1. The EMBO Journal 24: 4166–4175. doi:10.1038/sj.emboj.7600877

Mikhailov, M. V., Proks, P., Ashcroft, F. M., Ashcroft, S. J., 1998. Expression of functionally active ATP-sensitive K-channels in insect cells using baculovirus. FEBS Letters 429: 390–394.

Murshudov, G. N., 2016. Refinement of Atomic Structures Against cryo-EM Maps. Meth. Enzymol. 579: 277–305. doi:10.1016/bs.mie.2016.05.033

Nelson, P. T., Jicha, G. A., Wang, W.-X., Ighodaro, E., Artiushin, S., Nichols, C. G., Fardo, D. W., 2015. ABCC9/SUR2 in the brain: Implications for hippocampal sclerosis of aging and a potential therapeutic target. Ageing Res. Rev. 24: 111–125. doi:10.1016/j.arr.2015.07.007

Nichols, C. G., 2016. Adenosine Triphosphate-Sensitive Potassium Currents in Heart Disease and Cardioprotection. Card Electrophysiol Clin 8: 323–335. doi:10.1016/j.ccep.2016.01.005

Nichols, C. G., 2006. KATP channels as molecular sensors of cellular metabolism. Nature 440: 470–476. doi:10.1038/nature04711

Nichols, C. G., Shyng, S. L., Nestorowicz, A., Glaser, B., Clement, J. P., Gonzalez, G., Aguilar-Bryan, L., Permutt, M. A., Bryan, J., 1996. Adenosine Diphosphate as an Intracellular Regulator of Insulin Secretion. Science 272: 1785–1787. doi:10.1126/science.272.5269.1785

Noma, A., 1983. ATP-regulated K^+^ channels in cardiac muscle. Nature 305: 147–148. doi:10.1113/jphysiol.1993.sp019620/pdf

Pettersen, E. F., Goddard, T. D., Huang, C. C., Couch, G. S., Greenblatt, D. M., Meng, E. C., Ferrin, T. E., 2004. UCSF chimera - A visualization system for exploratory research and analysis. J Comput Chem 25: 1605–1612. doi:10.1002/jcc.20084

Pinney, S. E., MacMullen, C., Becker, S., Lin, Y.-W., Hanna, C., Thornton, P., Ganguly, A., Shyng, S.-L., Stanley, C. A., 2008. Clinical characteristics and biochemical mechanisms of congenital hyperinsulinism associated with dominant KATP channel mutations. J. Clin. Invest. 118: 2877–2886. doi:10.1172/JCI35414

Pratt, E. B., Zhou, Q., Gay, J. W., Shyng, S.-L., 2012. Engineered interaction between SUR1 and Kir6.2 that enhances ATP sensitivity in KATP channels. J Gen Physiol 140: 175–187. doi:10.1085/jgp.201210803

Proks, P., Antcliff, J. F., Lippiat, J., Gloyn, A. L., Hattersley, A. T., Ashcroft, F. M., 2004. Molecular basis of Kir6.2 mutations associated with neonatal diabetes or neonatal diabetes plus neurological features. Proceedings of the National Academy of Sciences 101: 17539–17544. doi:10.1073/pnas.0404756101

Proks, P., de Wet, H., Ashcroft, F. M., 2010. Activation of the K ATPchannel by Mg-nucleotide interaction with SUR1. J Gen Physiol 136: 389–405. doi:10.1085/jgp.201010475

Punjani, A., Rubinstein, J. L., Fleet, D. J., Brubaker, M. A., 2017. cryoSPARC: algorithms for rapid unsupervised cryo-EM structure determination. Nat. Methods 14: 290–296. doi:10.1038/nmeth.4169

Rohou, A., Grigorieff, N., 2015. CTFFIND4: Fast and accurate defocus estimation from electron micrographs. J. Struct. Biol. 192: 216–221. doi:10.1016/j.jsb.2015.08.008

Rorsman, P., Trube, G., 1985. Glucose dependent K^+^-channels in pancreatic beta-cells are regulated by intracellular ATP. Pflugers Arch. 405: 305–309. doi:10.1007/BF00581502.pdf

Rubaiy, H. N., 2016. The therapeutic agents that target ATP-sensitive potassium channels. Acta Pharm 66: 23–34. doi:10.1515/acph-2016-0006

Saint-Martin, C., Arnoux, J.-B., de Lonlay, P., Bellanné-Chantelot, C., 2011. KATP channel mutations in congenital hyperinsulinism. Seminars in Pediatric Surgery 20: 18–22. doi:10.1053/j.sempedsurg.2010.10.012

Scharschmidt, B. F., Keeffe, E. B., Blankenship, N. M., Ockner, R. K., 1979. Validation of a recording spectrophotometric method for measurement of membrane-associated Mg-and NaK-ATPase activity.J. Lab. Clin. Med. 93: 790–799.

Scheres, S. H. W., 2016. Processing of Structurally Heterogeneous Cryo-EM Data in RELION. Meth. Enzymol. 579: 125–157. doi:10.1016/bs.mie.2016.04.012

Scheres, S. H. W., 2012. RELION: Implementation of a Bayesian approach to cryo-EM structure determination. J. Struct. Biol. 180: 519–530. doi:10.1016/j.jsb.2012.09.006

Schwanstecher, C., Dickel, C., Panten, U., 1994. Interaction of tolbutamide and cytosolic nucleotides in controlling the ATP-sensitive K^+^ channel in mouse beta-cells. Br. J. Pharmacol. 111: 302–310. doi:10.1111/(ISSN)1476-5381

Schwappach, B., Zerangue, N., Jan, Y. N., Jan, L. Y., 2000. Molecular Basis for K ATP Assembly. Neuron 26: 155–167. doi:10.1016/S0896-6273(00)81146-0

Sharma, N., Crane, A., Gonzalez, G., Bryan, J., Aguilar-Bryan, L., 2000. Familial hyperinsulinism and pancreatic beta-cell ATP-sensitive potassium channels. Kidney Int. 57: 803–808. doi:10.1046/j.1523-1755.2000.00918.x

Shorter, K., Farjo, N. P., Picksley, S. M., Randall, V. A., 2008. Human hair follicles contain two forms of ATP-sensitive potassium channels, only one of which is sensitive to minoxidil. FASEBJ. 22: 1725–1736. doi:10.1096/fj.07-099424

Shyng, S., Nichols, C. G., 1997. Octameric stoichiometry of the KATP channel complex. J Gen Physiol 110: 655–664. doi:10.1085/jgp.110.6.655

Shyng, S. L., Nichols, C. G., 1998. Membrane phospholipid control of nucleotide sensitivity of KATP channels. Science 282: 1138–1141.

Spruce, A. E., Standen, N. B., Stanfield, P. R., 1985. Voltage-dependent ATP-sensitive potassium channels of skeletal muscle membrane. Nature 316: 736–738.

Standen, N. B., Quayle, J. M., Davies, N. W., Brayden, J. E., Huang, Y., Nelson, M. T., 1989. Hyperpolarizing vasodilators activate ATP-sensitive K^+^ channels in arterial smooth muscle. Science 245: 177–180.

Stanley, C. A., Thornton, P. S., Ganguly, A., MacMullen, C., Underwood, P., Bhatia, P., Steinkrauss, L., Wanner, L., Kaye, R., Ruchelli, E., Suchi, M., Adzick, N. S., 2004. Preoperative evaluation of infants with focal or diffuse congenital hyperinsulinism by intravenous acute insulin response tests and selective pancreatic arterial calcium stimulation. J. Clin. Endocrinol. Metab. 89: 288–296. doi:10.1210/jc.2003-030965

Stein, N., 2008. CHAINSAW: a program for mutating pdb files used as templates in molecular replacement. J Appl Crystallogr 41: 641–643. doi:10.1107/S0021889808006985

Sturgess, N. C., Ashford, M. L., Cook, D. L., Hales, C. N., 1985. The sulphonylurea receptor may be an ATP-sensitive potassium channel. Lancet 2: 474–475.

Tang, G., Peng, L., Baldwin, P. R., Mann, D. S., Jiang, W., Rees, I., Ludtke, S. J., 2007. EMAN2: an extensible image processing suite for electron microscopy. J. Struct. Biol. 157: 38–46. doi:10.1016/j.jsb.2006.05.009

Tantama, M., Martinez-Franois, J. R., Mongeon, R., Yellen, G., 2013. Imaging energy status in live cells with a fluorescent biosensor of the intracellular ATP-to-ADP ratio. Nat Commun 4: 2550. doi:10.1038/ncomms3550

Tao, X., Avalos, J. L., Chen, J., MacKinnon, R., 2009. Crystal Structure of the Eukaryotic Strong Inward-Rectifier K^+^ Channel Kir2.2 at 3.1 angstrom Resolution. Science 326: 1668–1674. doi:10.1126/science.1180310

Tucker, S. J., Gribble, F. M., Proks, P., Trapp, S., Ryder, T. J., Haug, T., Reimann, F., Ashcroft, F. M., 1998. Molecular determinants of KATP channel inhibition by ATP. The EMBO Journal 17: 3290–3296. doi:10.1093/emboj/17.12.3290

Tucker, S. J., Gribble, F. M., Zhao, C., Trapp, S., Ashcroft, F. M., 1997. Truncation of Kir6.2 produces ATP-sensitive K^+^ channels in the absence of the sulphonylurea receptor. Nature 387: 179–183. doi:10.1038/387179a0

Whorton, M. R., MacKinnon, R., 2011. Crystal structure of the mammalian GIRK2 K^+^ channel and gating regulation by G proteins, PIP_2_, and sodium. Cell 147: 199–208. doi:10.1016/j.cell.2011.07.046

Zhang, Z., Chen, J., 2016. Atomic Structure of the Cystic Fibrosis Transmembrane Conductance Regulator. Cell 167: 1586–1597.e9. doi:10.1016/j.cell.2016.11.014

Zheng, S. Q., Palovcak, E., Armache, J.-P., Verba, K. A., Cheng, Y., Agard, D. A., 2017. MotionCor2: anisotropic correction of beam-induced motion for improved cryo-electron microscopy. Nat. Methods 14: 331–332. doi:10.1038/nmeth.4193

Zhou, M., Li, Y., Hu, Q., Bai, X.-C., Huang, W., Yan, C., Scheres, S. H.W., Shi, Y., 2015. Atomic structure of the apoptosome: mechanism of cytochrome c-and dATP-mediated activation of Apaf-1. Genes Dev. 29: 2349–2361. doi:10.1101/gad.272278.115

